# The SWAN atlas: an open multimodal resource combining neuroimaging and high-resolution histology of the human brain in stereotactic space

**DOI:** 10.64898/2026.10.12.758634

**Authors:** Eduardo Joaquim Lopes Alho, Helen Friedrich, Ana Tereza Di Lorenzo Alho, Armando Alaminos Bouza, Edson Amaro, Lea Tenenholz Grinberg, Simon Oxenford, Erich Talamoni Fonoff, Andreas Horn, Helmut Heinsen

**Affiliations:** Morphological Brain Research Unit, Department of Psychiatry, University of Würzburg, Germany; Division of Functional Neurosurgery, Department of Neurology, University of São Paulo Medical School, São Paulo, Brazil; Department of Radiology, Hospital das Clínicas HCFMUSP, Faculdade de Medicina, Universidade de São Paulo, São Paulo, Brazil; Department of Neurology, Neuromodulation and Movement Disorders Unit, Charité – University Medicine (CCM), Berlin, Germany; Memory and Aging Center, Department of Neurology, Weill Institute for Neurosciences, University of California, San Francisco, California, USA; Center for Brain Circuit Therapeutics, Department of Neurology, Brigham & Women’s Hospital, Harvard Medical School, Boston, MA, USA; Julius-Maximilian-University Würzburg, Germany; Network Stimulation Institute, Department of Stereotactic and Functional Neurosurgery, University Hospital Cologne, Cologne, Germany; Clinica de Dor e Funcional (CDF), São Paulo, Brazil; Mevis Informática Médica LTDA, São Paulo, Brazil

**Keywords:** human brain atlas, human histology, stereotactic surgery, deep brain stimulation, functional neurosurgery, MNI space, neuroanatomy

## Abstract

Precise localization of human brain regions is critical in clinical practice and population neuroimaging, particularly in stereotactic functional neurosurgery, where small targeting errors may lead to treatment failure. Although MRI provides *in vivo* and *postmortem* brain parcellation, histology offers complementary mesoscopic and microscopic information. A major limitation of histological atlases for clinical use is accurate coregistration with *in vivo* imaging, since tissue processing introduces deformation. To reduce this problem, we sectioned an entire human brain using high-thickness slices, which are less prone to slice-level deformation. Histological sections were registered to the brain’s own MRI and normalized to standard stereotactic space through a multistep transform chain. Users can apply MRI-to-MRI registration to transform the atlas into patient space and vice versa, with further nonlinear refinement using SwanWarp or WarpDrive tools. Segmentations and software are being prepared for open distribution. This resource may empower both clinical neurosurgical research and the broader neuroimaging field at large.

**Resource scope:** SWAN is designed as a free, practical and reproducible bridge between high-resolution human histology and patient-oriented stereotactic neuroimaging. The resource combines source data, transform chains, segmentations, and 3D Slicer- and Lead-DBS compatible workflows that allow investigators, clinicians and trainees to use the default atlas or create their own anatomical labels, inspect patient-relevant regions of interest against histology and support research, planning and post hoc analysis in functional neurosurgery, neurology and translational neuroimaging. The SWAN Atlas Viewer also enables users to interactively navigate across three-dimensional histological and imaging spaces, providing a powerful educational tool for understanding spatial relationships between tissue anatomy and neuroimaging. The SWAN project website (www.swanatlas.com) serves as the central public entry point for the resource, providing project documentation, representative histological and multimodal atlas content, software information, and information on the planned data and code repositories.

## Introduction

In recent decades, magnetic resonance imaging (MRI) has developed extensively, providing high quality brain images. MRI may reveal structural and functional images noninvasively, through development of diverse pulse sequences and image processing algorithms. At times, even detailed tissue properties such as iron content and water diffusion can be quantified *in vivo,* rendering MRI the method of choice to study individual brain anatomy noninvasively. Although application of high magnetic fields and development of dedicated sequences have facilitated to reconstruct images with high resolution and contrast^1^, some properties of the complex neural tissue may still be better visualized based on histological staining of brain slices^2^.

This is of relevance in functional stereotactic neurosurgery where small brain structures are routinely targeted for treatment of brain disorders. Deep brain stimulation (DBS) electrodes rely on precise guidance to anatomical targets in order to achieve optimal clinical results^3^. Understanding the three-dimensional relationships of DBS electrode contacts and their electric fields relative to the surrounding anatomy is key to maximize outcomes and avoid undesired effects. Several human brain atlases have been previously proposed^4–24^, and animal atlases have been published in rodents^25–31^ and non-human primates^32–38^. According to a comprehensive review on brain atlases^39^, there is a resurgence of interest on the topic motivated by functional imaging techniques that yield results of higher resolution. Hence, brain atlases are not only relevant during DBS surgery and postoperative titration of electrical current, but they might be helpful for proposing new stereotactic targets and optimizing existing ones.

The FOCUS atlas recently extended subcortical anatomical mapping by integrating ultra-high-resolution ex vivo MRI with histology, fibre dissections and neuroanatomical references to define deformable grey- and white-matter structures at 100 µm isotropic resolution^40^. While FOCUS emphasizes a detailed synthesis of subcortical structures and pathways for stereotactic applications, SWAN provides complementary access to serial whole-brain histology from a single specimen, its blockface images and *in situ* MRI, together with forward and inverse transformations linking these source modalities to stereotactic space. This framework allows users to inspect the histological basis of an anatomical label, revise its boundaries or delineate additional structures and propagate those changes across spaces.

Histological atlases in book format have played a major role in the history of functional neurosurgery. Influential atlases such as the Schaltenbrand-Wahren^7^ or Talairach atlases^13^ have been critical stepping stones to develop precision neurosurgery. Later, computational resources have become available to digitize atlases and register them to stacks as 3D volumes that may be appreciated within dedicated software^41^. While promising and theoretically very useful, thus far, the process has suffered from the key issue that thin slices are nonlinearly deformed when scanned under the microscope. This leads to two drawbacks: First, the registration of slices to volume format is inherently complicated since each slice is deformed in different ways. Second, these nonlinear deformations complicate precise co-registration with *in-vivo* imaging ^42–45^. In order to reduce the impact of some of these shortcomings, in 2017, our group proposed the use of high-thickness histological sections with a modified Nissl stain^46,47^. Thicker slices are physically less prone to nonlinear distortions and may therefore be more accurately registered to one another. Furthermore, our slices have been precisely co-registered to an *in-situ* MRI of the same brain (obtained before brain extraction from the skull within a short post-mortem interval of ∼14 h)^48^. With this methodology, we were able to minimize the 2D and 3D distortions of histological processing and thus to enhance contrast between nuclear boundaries. Co-registration of histology to its own MRI and normalization to MNI space rendered a coherent histological atlas available in 3D^48^.

In the present resource paper, we characterize this whole human brain dataset cut in serial sections of 430µm thickness. The São Paulo Würzburg Atlas of Neuroanatomy (SWAN) dataset includes darkfield illumination images focusing on the subcortex, *in situ* 3T MRI including DTI, blockface 3D block and high-resolution whole brain histology, all normalized to MNI standard space. The transform chain is stored as forward and inverse transforms and is intended for distribution through an open-source 3D Slicer module and Lead-DBS-compatible workflows. Expert-segmentation labels in blockface space and in standard stereotactic (MNI) space have also been generated.

The SWAN project website (https://www.swanatlas.com) serves as the central public entry point for project documentation, representative histological and multimodal atlas content, software information and the staged release plan. Data and code are being prepared for deposition in versioned public repositories, as detailed in the Data availability and Code availability sections.

The workflow providing schematic overview of the atlas construction is shown in figures 1 and 2.

**Figure 1.**
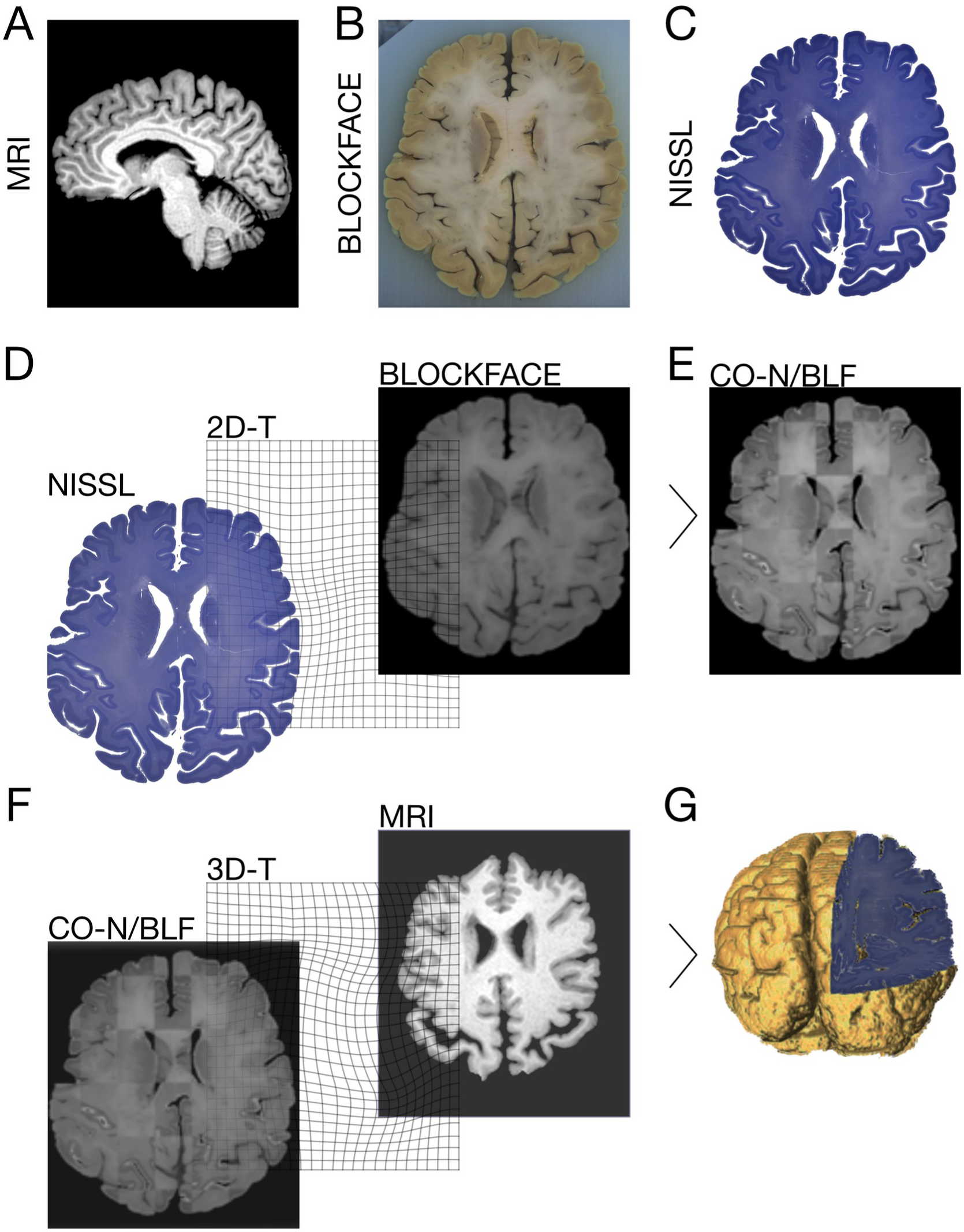
Workflow with schematic overview of the histological processing from **A** MRI scanning until reconstruction of **G** a coherent 3D histological dataset. Bidimensional (2D-T) and three-dimensional (3D-T) transforms are stored for distribution for further application in histological masks created in high resolution images. Abbreviations: CO-N/BLF = coregistered Nissl- and Blockface dataset.

**Figure 2.**
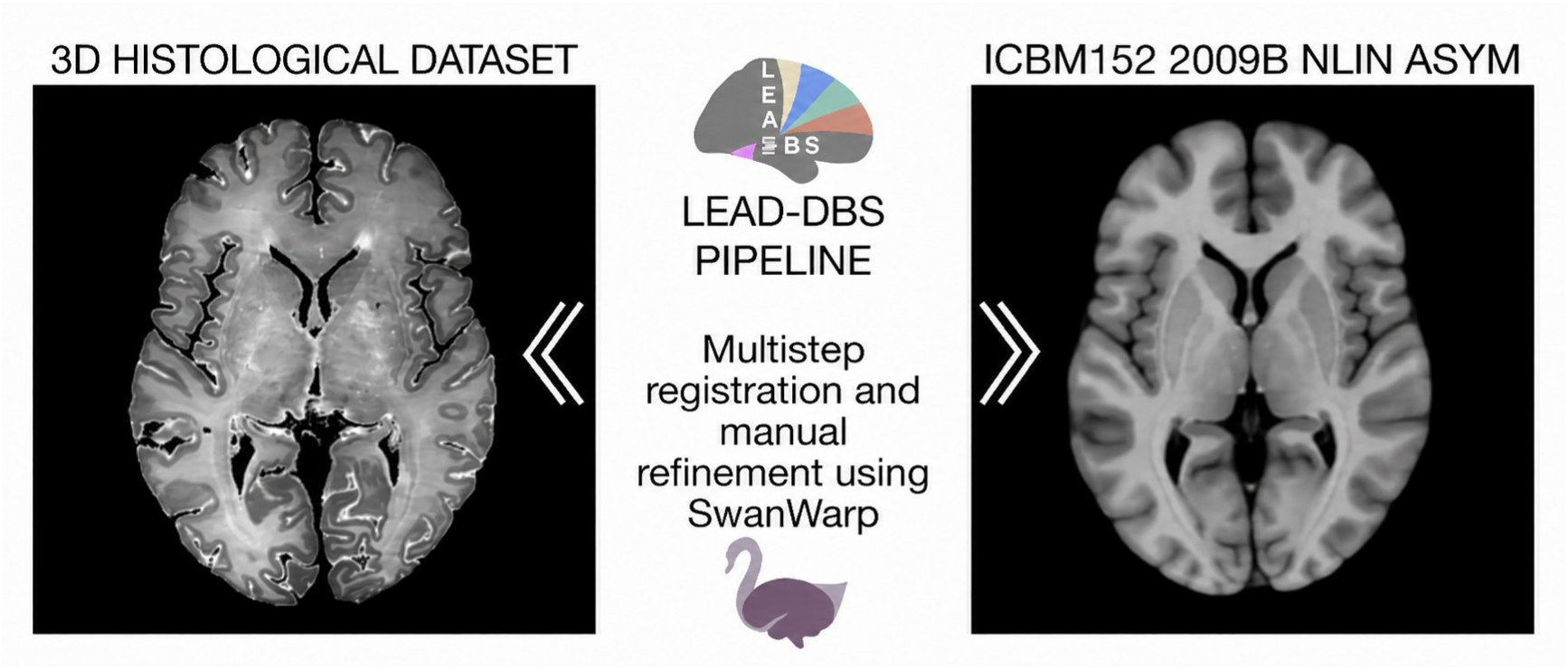
Registration of the histological volume with the ICBM 152 2009b Nlin Asym (MNI) space.

## Methods

The dataset comprises the brain of a woman who died at an age of 66 years from an acute myocardial infarct. Members of the family who agreed to participate in an elaborate post-mortem interview^49^ did not report neurological symptoms or deficits. Written informed consent from the family was obtained as part of a research protocol approved by the ethics board at University of São Paulo (project number 0086/09, approved by the National Commission of Ethics in Research -CONEP-under number 16029). After the family’s agreement, images from 3T MRI (Achieva, Philips Medical Systems, Netherlands) equipped with an eight-channel head coil, and 80 mT/m gradients were obtained prior to autopsy. MRI sequences were as follows: T1: TR = 6.3 ms, TE = 2.9 ms, TI = 791 ms, 1 x 1 x 1 mm voxels 3D; Axial FLAIR: TR=8613ms, TI = 2800 ms, TE = 130 ms, 0.65 mm x 0.87 mm, 1 NEX 3 mm/0.3 mm, FOV 230 mm x 183 mm, 2D, DTI: b = 3000, TE = 65 ms, TR 23.650 ms, FOV 256 mm x 256 mm, resolution 2.0 mm x 2.0 mm x 2.0 mm, NEX 2, gradient 32 directions, SENSE = 2 (AP), Flip angle = 90. An extensive infarct region mainly comprising the deep white matter and of the left parietooccipital lobes together with smaller liquefied cortical infarcts were diagnosed in 3T scans and confirmed by pale subcortical tissue after gallocyanin staining. In addition, small bilateral foci of neuron loss were detected in 430 µm thick gallocyanin-stained serial sections, mainly in the pre- and postcentral gyri. Some of these foci were closely associated with penetrating cortical vessels, others were confined to a laminar pattern affecting layer III and V of the pre- and postcentral regions. These foci with distinct neuron loss without central liquefying necroses could not be identified on 3T images.

### Histological procedure

After returning from radiology, the brain was extracted from the skull with a post-mortem interval of 13 hours and 5 minutes, and prepared for histological processing as described by Heinsen et al. in 2000^47^. Briefly, the brain was fixed by immersion in 20% formalin for 4 months. Formalin was changed three times in the first 3 weeks. After this period, the brain was dehydrated in graded series of ethanol solutions (70, 80 and 96%), for 1 week per stage, and the tissue was hardened by this procedure. After the final dehydration, the brain was placed into a transparent plastic box, filled with 8% celloidin (180-200 g of commercial alcohol-moistened celloidin dissolved in a mixture of 1000 ml 100% ethanol and 1000 ml diethyl ether). This custom-made box was placed into a desiccator, and the air was evacuated by a membrane vacuum pump (vacuum -150 mb from ambient air pressure) for 48 h. With this procedure, the celloidin was forced to enter the depth of the sulci and the ventricular system while being concentrated to 16%. However, celloidin did not infiltrate the brain. The plastic box was then placed into an aquarium, together with a petri dish (185 mm diameter) filled with chloroform. The celloidin was further hardened by chloroform vapors. The aquarium was opened daily to remove excess of ether/alcohol from the block’s surface. After 15 days, the celloidin became opaque and acquired a consistency of India rubber, retracting from the walls of the plastic box. Final hardening with 70% ethanol was achieved in 2 days. The celloidin block was trimmed to fit the size of the microtome, and glued onto the microtome with 8% celloidin, placed with 70% ethanol for 4 h to harden the glue. The block was serially sectioned on a sliding microtome (Polycut, Cambridge Instruments, UK) with 430 µm thickness, in the axial plane. Blockface imaging was obtained with a single lens reflex camera, set in a fixed position relative to the block’s surface. The slices were then stained according to the procedures described by Heinsen and Heinsen in 1991^46^. The sections were left overnight in petri dishes with a solution of performic acid (1 part of 100% formic acid, 3 parts of 30% H_2_O_2_ and 6 parts of distilled water), and constantly stirred on a electromechanical shaker with filter paper on the surface of the performic acid to keep the sections immersed in fluid. The acid was removed the next morning by running tap water for 1 h. The sections were stained in gallocyanin-chromalum with constant slight stirring for 3 h, dissolving 1.5g gallocyanin (Aldrich-Chemie, Steinheim, FRG) in 1000 ml of 5% potassium chromium (III) sulfate dodecahydrate (Merck). This mixture was boiled for 10 minutes, and after cooling and filtering, distilled water was added to restitute the volume of the staining solution up to 1000 ml. The resulting pH of the solution was 2.0. Gallocyanin excess was removed by thorough rinsing in running water for 1 h. The sections were then transferred from tap water into 50% ethanol and flattened by squeezing them between 2 sheets of filter paper. Thereafter, individual sections were pressed between filter paper and stored in stacks of perforated plates of stainless steel (170×120×1.5mm). This procedure was followed by dehydration in graded series of alcohol (1 h in 80% ethanol, overnight in propanol 100%, 2h in xylene 1:1, fresh xylene overnight. The slices were immersed in Permount (Fisher Scientific, Pittsburgh, PA) for 1 minute, and mounted (resin Permount) onto custom size (for whole brain) glass slides for microscopy. Lead blocks were placed on the coverslide for 2 weeks to prevent formation of air bubbles and keep the mounted sections flat. The complete polymerization of Permount that seals thick sections is a long-lasting process that comprises years. In general, stained thick sections should always be kept horizontally. Polymerization proceeds in a centripetal manner. After 4 weeks, the polymerized periphery of the coverslipped Permount ‘lake’ will prevent leakage of its central parts and the stained sections were photographed on a Kaiser LED light box with a Nikon D800E SLR digital camera mounted with a 50mm Sigma Makro lens. All photographs were acquired at a fixed distance between the camera sensor and the focused slides, ensuring a constant reproduction scale across the complete set of stained serial sections. Calibration rulers created with drawing software were printed on transparent folia, placed at the periphery of the slides. They served to calculate the reproduction scale and to transform the calibrated distances into pixels. The whole data set spanned 279 images, each with 0.0250 x0.0250 x 0.430 mm voxel resolution, stored in JPEG format.

Dark field illumination photographs were obtained with a smaller field of view for brainstem, subthalamic region and thalamus.

**Figure 3.**
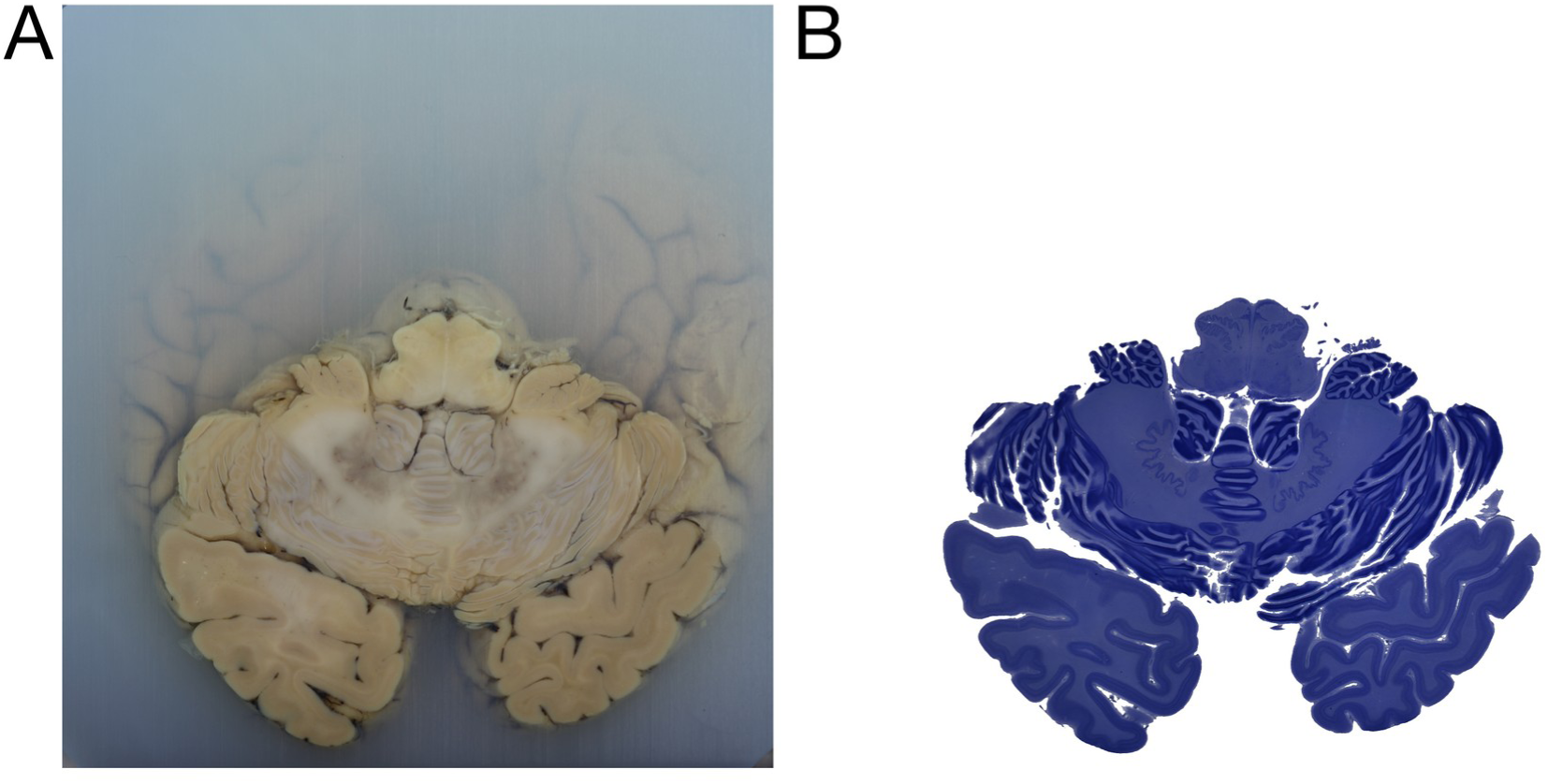
In (A), the celloidin blockface and in (B) the Nissl-stained slice from the surface of the block in (A)

### Image preprocessing and registration procedures

As the histological processing causes inevitable case-specific three-dimensional and two-dimensional deformations of processed brain tissue, a computational pipeline was developed by our group in order to restore the final histology sections to their original form (*in situ* MRI)^50^.

### Bidimensional registration of volumes

Initially, both blockface and histological images were segmented from the background, to eliminate unnecessary information. To achieve this, we used an orthogonal color model that allows pixels to cluster in specific regions in color space^51^. The images were transformed from RGB to YIQ space. The tissue pixels were modeled as a Gaussian mixture model. The pixels were classified using this model, generating an initial binary mask, further refined by active contours. After segmenting both blockface images and histological slices, they were converted to grayscale. This segmentation method was implemented in MATLAB and is described in more detail elsewhere^50^. The next step for MRI-histology fusion was to perform 2D registration between blockface images with their corresponding histological plates (2D T). This step was achieved using the affine registration method implemented in Advanced Normalization Tools (ANTs; http://stnava.github.io/ANTs/; ^52^) using Mattes mutual information as similarity metric^53^.

Original slice numbering was preserved throughout the 2D stage, with unavailable histological section 147 excluded from registration and section 001 excluded because of insufficient usable tissue signal. For each available section, the light-field image was first cropped using the same horizontal range used during volume construction (x = 850–6000 in MATLAB indexing) and converted to grayscale. Each cropped section was then registered to the corresponding grayscale blockface image using an affine 2D registration initialized from tissue masks and optimized with mutual information. Slices with residual misalignment after automatic registration were manually refined using paired anatomical landmarks placed in Fiji; these landmarks were used to estimate an incremental 2D affine correction from the automatically warped histological image to the blockface reference. The final per-slice transform was defined as the composition of the automatic affine transform and the approved manual correction when present. The same final per-slice transforms were then applied to the corresponding grayscale images, histological RGB images and their 2D registered darkfield pairs. Final 2D outputs were retained as PNG image stacks for visual quality control prior to volumetric reconstruction.

Registration quality was assessed visually using red-green overlays and quantitatively using tissue-mask Dice coefficient, Jaccard index, centroid distance and mutual information before and after registration. Normalized cross-correlation was not used as a primary metric because blockface and histological images have different intensity profiles and contrast mechanisms. Per-slice metrics were exported to tabular format, and representative slices were selected across the rostrocaudal extent of the volume to generate summary figures combining qualitative overlays and quantitative registration metrics.

**Figure 4:**
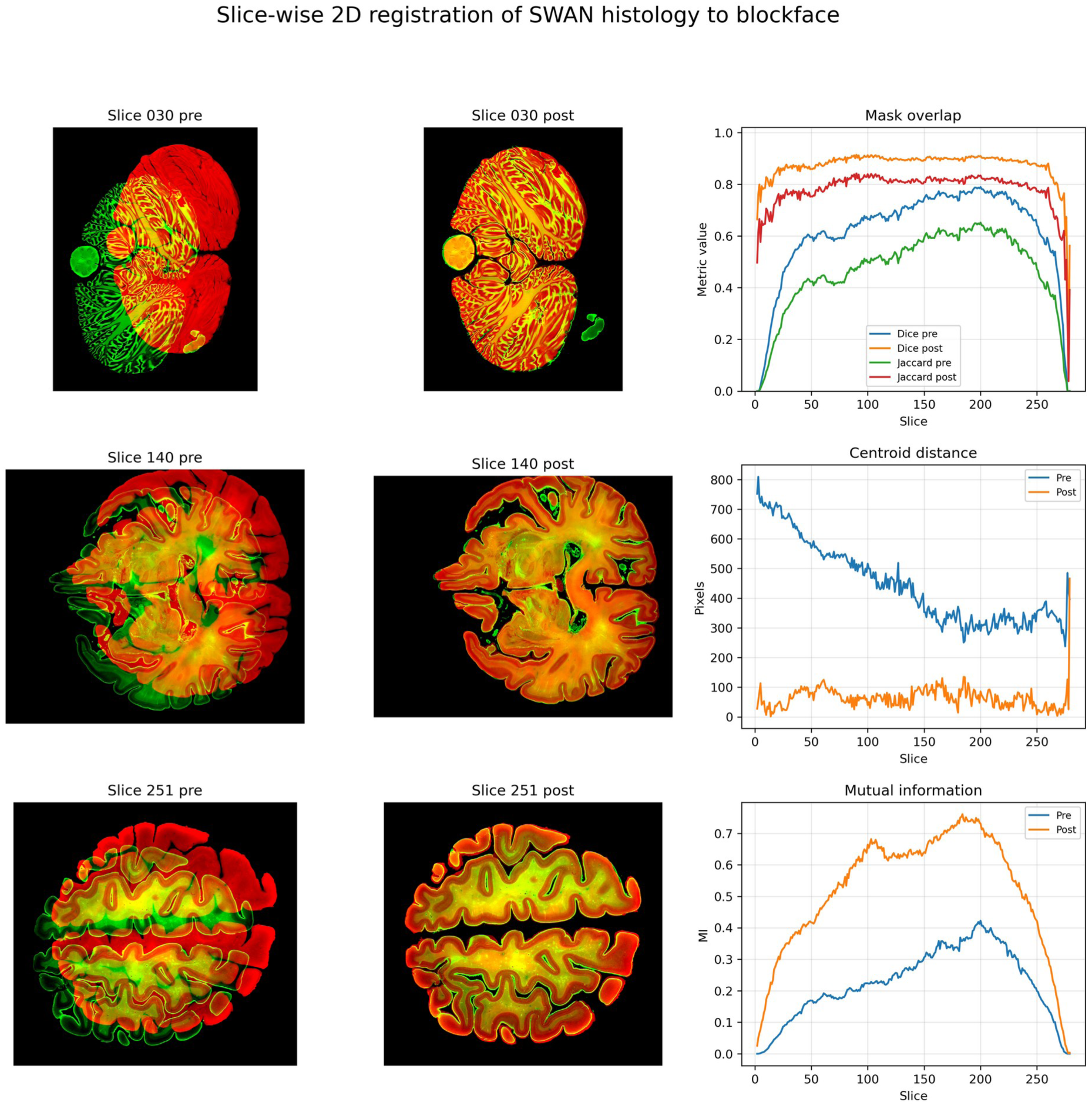
Automatic slice-wise 2D registration of SWAN histology to blockface images. Representative sections illustrate alignment before and after automatic registration. Registration performance is summarized using Dice and Jaccard overlap, centroid distance, and mutual information. These data precede manual landmark-based refinement in Fiji; final post-refinement metrics are reported in the Results.

The images were then stacked onto an intermediary 3D volume, stored in NIfTI format. This 3D volume was roughly aligned to the MRI manually to create an initial overlap between the datasets, allowing the registration algorithms to work properly.

### 3D MRI histology registration and SwanWarp refinement

Preprocessing of the *in situ* MRI allowed better MRI-histology registration. Field inhomogeneity correction with N4 algorithm^54^, skull stripping with FSL’s brain extraction tool (BET) algorithm (FMRIB Software Library v6.0, Analysis Group, Oxford UK) and cubic spline sampling (implemented in ITK) were applied in the volumetric T1 images.

Three-dimensional MRI-histology registration (3D T) was achieved using nonlinear symmetric diffeomorphic method (SyN)^55^, and saved for later use. The histology-derived volume was manually refined into the subject MRI space using SwanWarp, a landmark-based volumetric registration self-developed tool, that allows interactive correction of histology-to-MRI alignment, or any 3D volumes. SwanWarp is developed as a module for 3D Slicer software (https://www.slicer.org/) and is being developed as part of the SwanAtlas Pipeline. For users, it will be useful to precisely refine warps between SWAN histological labels into patient space.

The brain MRI volume was used as the fixed image, the histological volume as the moving image, and corresponding anatomical landmarks were placed as paired moving/fixed points across multiple planes. Additional anchor points were used as identity constraints to stabilize regions that did not require deformation. A deformable grid transform was then estimated from the paired landmarks and anchors using Plastimatch within 3D Slicer, applied to the histological volume and hardened to generate the final Histology-in-MRI volume. This transform was applied to all previously registered volumes (blockface, darkfield and light field), so all MRI sequences of the subject were registered to their corresponding histological datasets.

Registration quality for the SwanWarp refinement was assessed using the original histological volume and the warped histological volume resampled to the fixed MRI grid. Automatic quality-control metrics included robust intensity normalization, mutual information, normalized mutual information, normalized cross-correlation, edge-map correlation, automatic edge Dice and automatic edge Jaccard. Multiplanar overlay, checkerboard and edge-overlay images were generated for qualitative inspection. Forward-inverse consistency was evaluated by applying the forward SwanWarp transform followed by its inverse to sampled points and measuring round-trip displacement error in millimeters.

### 3D MRI histology to MNI space normalization

The registration of this histological dataset into MNI standard space (ICBM152 2009b Nlin Asym) was performed in a multistep registration procedure in the open source environment, Lead-DBS^3,56^. This was done using the “effective: low variance” preset with subcortical refinement that was implemented in ANTs and concluded with two consecutive SyN stages. This parameter set was top-performing for subcortical structures in a study that involved >11,000 nonlinear deformations using various tools^57^. In a recent second comparative study of nonlinear registrations performed by a different group, this approach was also top performer^58^.

This registration served as an – already precise – initial registration between the whole-brain histological atlas and the MNI space. It was further manually refined by an expert (H.F.) using a novel tool that allows for manual interaction with nonlinear warp fields. The method – termed “WarpDrive” exists within Lead-DBS and allows for small manual adjustments of nonlinear registrations with the mouse^59^. In meticulous work spanning over multiple sessions (total of ∼16 work hours), slight misalignments between histology and MNI template could thus be corrected (Figure 2).

Segmentation of several anatomical structures were performed in the original high-definition histological images and dark-field images in blockface space and then propagated to MNI space by applying the transform chain. This open-source pipeline allows any user to create their own segmentation in our histological dataset, for structures of particular interest, generating their own atlas.

**Figure 5.**
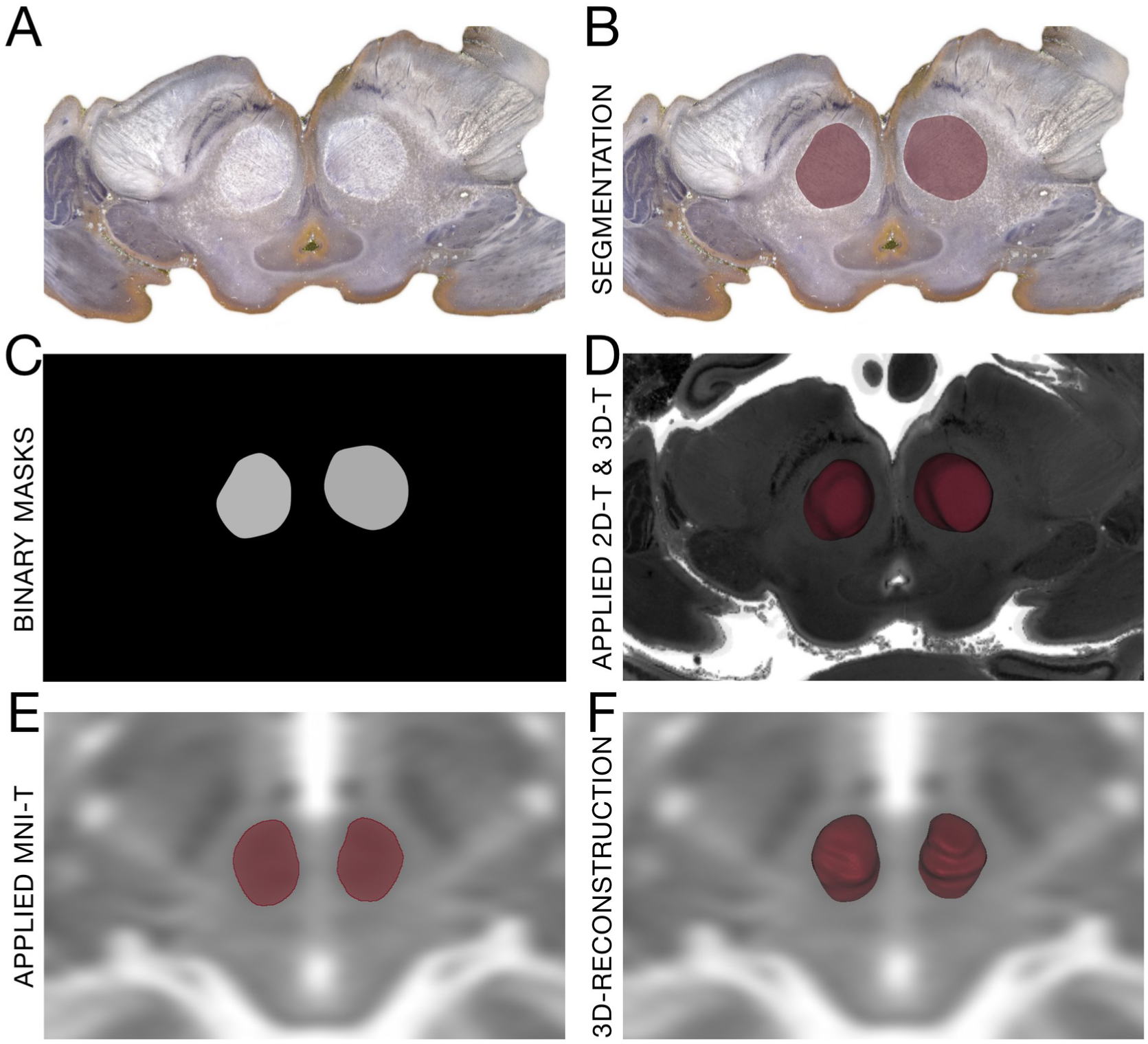
Creation of histological masks in high resolution histology and applying the transforms to combine the histological labels in MNI space.

### Inverse-chain diagnostics

Inverse label testing maps final-MNI labels back to blockface space to determine whether a segmentation in MNI space falls on the expected histological anatomy. A diagnostic low-resolution red nucleus inverse test used true numerical inversion for the transforms yielding Dice 0.909 against the original low-resolution blockface label.

## Results

### Registration metrics and technical validation

Registration performance was evaluated using complementary spatial and intensity-based metrics. The Dice coefficient quantifies the overlap between binary tissue masks and ranges from 0, indicating no overlap, to 1, indicating complete overlap. The Jaccard index similarly measures mask overlap, but is more stringent because it is defined as the intersection divided by the union of the two masks. Centroid distance measures the Euclidean distance, in pixels, between the centers of mass of the blockface and histological tissue masks, providing a simple estimate of residual global displacement after registration. Mutual information was used as an intensity-based similarity metric, which is particularly useful for multimodal registration because it does not require identical contrast between the two images. In the present dataset, the final slice-wise registration increased Dice and Jaccard overlap across most sections, markedly reduced centroid distance, and increased mutual information relative to the pre-registration condition. Together with the visual overlays, these findings indicate that the final 2D transformation chain improved both global tissue positioning and local anatomical correspondence between SWAN histology and blockface space. Because the blockface and SWAN images have different contrast mechanisms, the overlap-based metrics were interpreted together with visual inspection rather than as standalone measures of anatomical accuracy.

For 2D histology-to-blockface registration, perfect overlap is not expected because the two modalities differ in tissue contrast, illumination, section deformation, tears, missing fragments, and local histological distortions. In this context, Dice and Jaccard values in the high range after registration, together with low centroid displacement and consistent visual overlap of anatomical boundaries, indicate a strong registration result. The most relevant expectation is not mathematical identity between masks, but preservation of tissue topology and accurate alignment of major anatomical contours and internal landmarks. In the present dataset, the final registration achieved this pattern: overlap metrics improved across the series, centroid displacement was markedly reduced, and the post-registration overlays showed close correspondence of cortical boundaries, cerebellar folia, ventricular spaces, and major tissue fragments. These results support the use of the final slice-wise transforms for subsequent volumetric reconstruction and downstream mapping into blockface space.

The rebuilt registration quality-control package evaluated 277 registered sections using tissue-mask overlap, centroid-displacement and mutual-information metrics. Dice increased from 0.6164 ± 0.1816 before registration to 0.8720 ± 0.0720 after final registration, while Jaccard increased from 0.4661 ± 0.1592 to 0.7784 ± 0.0848. Centroid distance decreased from 445.26 ± 135.61 pixels to 63.30 ± 37.37 pixels, corresponding to an 84.3% mean reduction in global tissue-mask displacement. Mutual information increased from 0.2234 ± 0.1112 to 0.5181 ± 0.1903. All evaluated slices improved for Dice, Jaccard and mutual information, and 99.6% improved for centroid distance. Because blockface and histological images differ in contrast, illumination, local tissue deformation, tearing and missing fragments, overlap metrics were interpreted together with representative red-green overlays rather than as standalone measures of anatomical identity.

**Figure 6.**
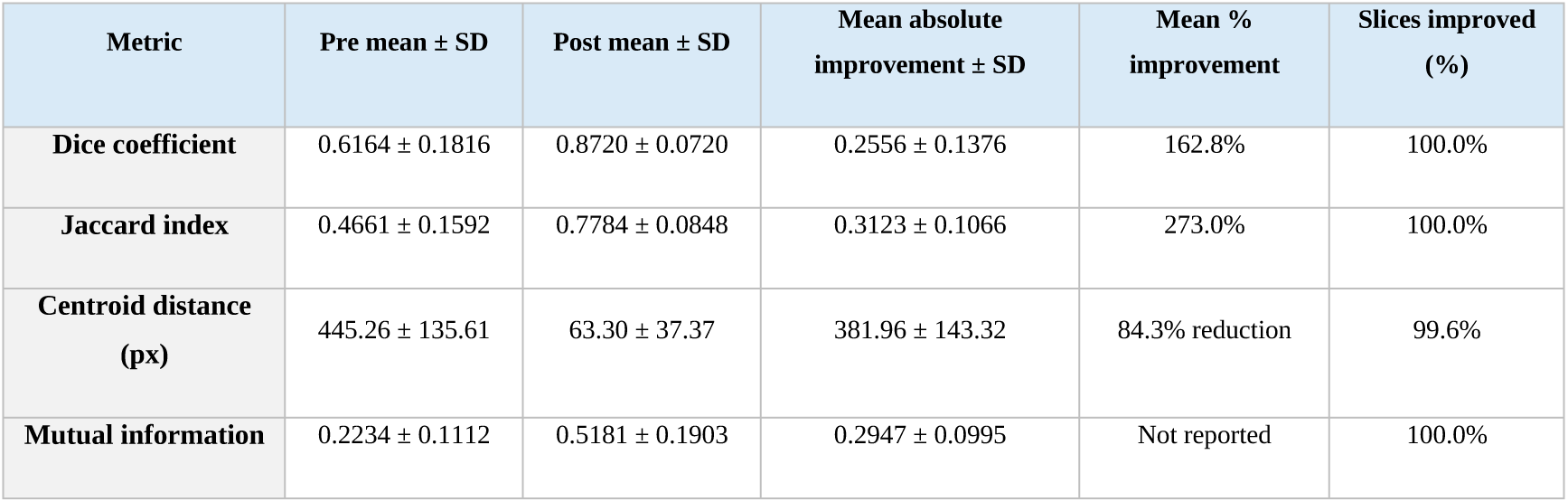
Registration performance was quantified across 277 registered sections using tissue-mask Dice coefficient, Jaccard index, centroid distance, and mutual information. Dice increased from 0.6164 ± 0.1816 before registration to 0.8720 ± 0.0720 after final registration, while Jaccard increased from 0.4661 ± 0.1592 to 0.7784 ± 0.0848. Centroid distance decreased from 445.26 ± 135.61 px to 63.30 ± 37.37 px. Mutual information increased from 0.2234 ± 0.1112 to 0.5181 ± 0.1903. These quantitative improvements, together with visual inspection of representative overlays, support accurate slice-wise alignment of SWAN histology to blockface space.

**Figure 7.**
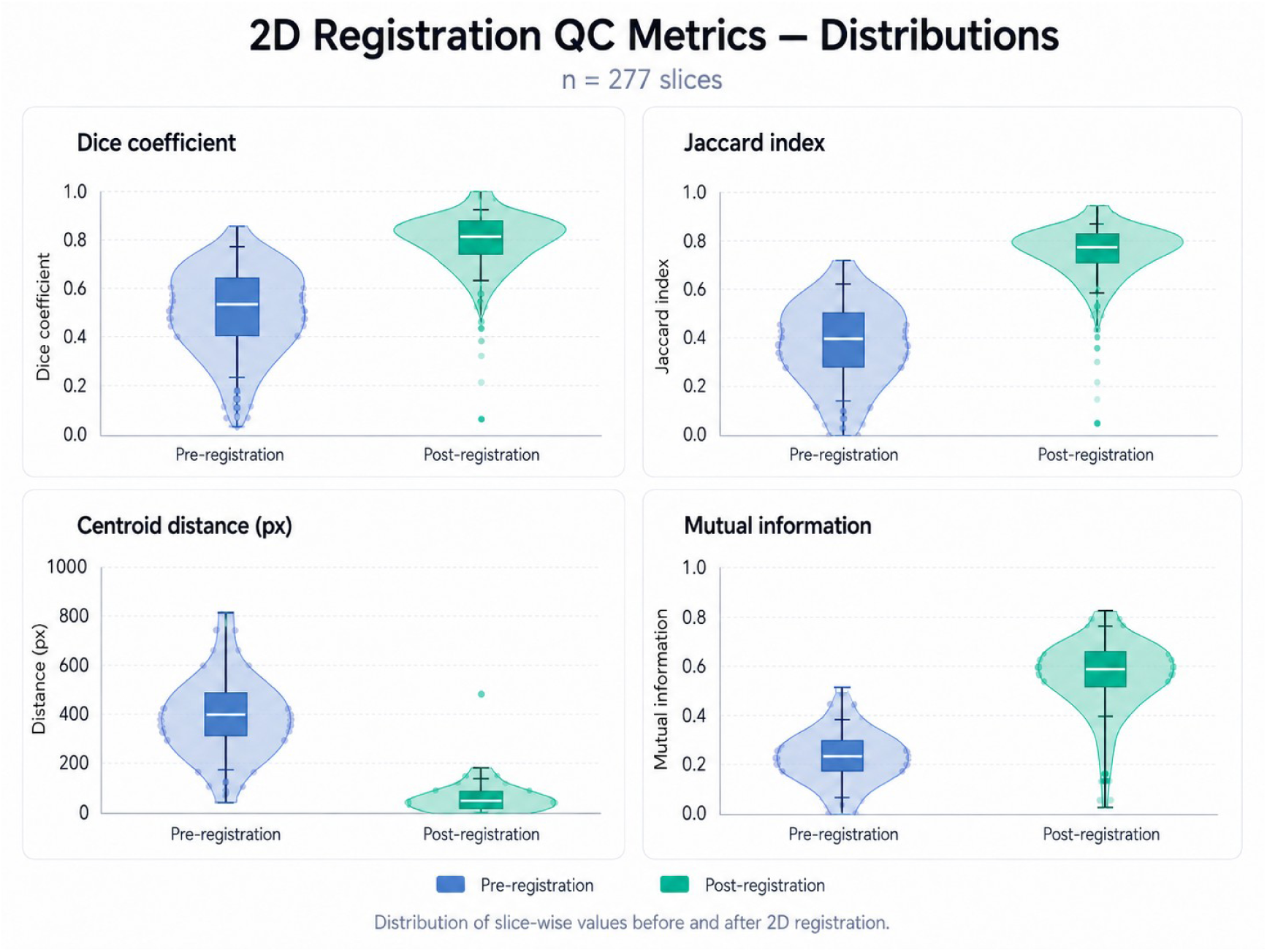
Distributions of 2D registration metrics. The 277 slices show improved overlap, reduced centroid distance and increased mutual information after registration.

### SwanWarp 3D refinement for localized corrections

SwanWarp refinement produced visually improved alignment between the histological volume and the subject MRI, particularly along cortical, ventricular and cerebellar boundaries, as assessed by multiplanar overlay and checkerboard inspection. Automatic image-based metrics showed largely stable global intensity similarity after warping, with mutual information changing from 0.12695 to 0.12494, normalized mutual information from 1.01995 to 1.01964, and normalized cross-correlation from 0.46755 to 0.46435. Edge-based agreement showed a modest improvement, with edge correlation increasing from 0.30669 to 0.31331, corresponding to a 2.16% relative increase. Forward-inverse consistency analysis demonstrated high numerical stability of the deformation field: across 500 sampled points, the mean round-trip error was 0.00119 mm, the median error was 0.000078 mm, the 95th percentile was 0.00702 mm, and the maximum error was 0.00993 mm.

Deformation-field analysis showed that SwanWarp refinement introduced spatially localized, moderate corrections while preserving the overall topology of the volume. Across 237,053 sampled voxels, the mean displacement magnitude was 0.720 mm and the median displacement was 0.426 mm. The interquartile range was 0.099–1.091 mm, and the 95th percentile displacement was 2.367 mm, with a maximum sampled displacement of 8.340 mm. Jacobian analysis demonstrated a deformation field centered close to local volume preservation, with a mean Jacobian determinant of 1.014, a median of 1.006, and a 5th–95th percentile range of 0.864–1.153. No non-positive Jacobian determinants were detected, supporting the absence of folding or local non-invertibility in the sampled deformation field.

### Segmentations

Table 1 (Supplementary Material) lists 127 anatomical structures and subdivisions, organized into 11 groups, and their abbreviations used in SWAN. The label set covers the basal ganglia and associated nuclei; basal forebrain cholinergic fields; amygdala and transition regions; hypothalamic and mammillary regions; thalamus and metathalamus; deep cerebellar nuclei; brainstem nuclei and regional complexes; pallidal pathways and the fields of Forel; brainstem and cerebellar pathways; limbic fibre bundles; and internal-capsule and commissural pathways. Whole structures and their subdivisions are identified separately in the table.

Anatomical labels were delineated in high-resolution histological and darkfield images in blockface space by a neuroanatomy expert. Regional boundaries were assessed against the tissue contrast and anatomical relationships visible in the corresponding sections, with published atlases and structure-specific studies providing nomenclature and anatomical guidance. Labels were then propagated through the documented transformation chain to the specimen MRI and final MNI space. The editable source segmentations retain the link to the underlying histology, enabling boundaries to be inspected and revised in their original anatomical context^48^.

Reference anatomy included the Julich-Brain cytoarchitectonic atlas and its version 3.1 maps^60,61^. Thalamic terminology and regional boundaries were informed by the Schaltenbrand–Wahren, Morel and Krauth atlases^7,15,16,23^. Structure-specific references included the basal forebrain maps of Zaborszky and colleagues^22^. Amygdalar subdivisions were guided by the cytoarchitectonic and multimodal studies of Amunts and Kedo and colleagues^62,63^. The dorsal BNST was interpreted with reference to the human study by Brandstetter and colleagues^64^.

Deep cerebellar nuclei were interpreted using the cytoarchitectonic descriptions of Tellmann and colleagues^65^. Brainstem anatomy was guided by established stereotactic descriptions^11^. Prior SWAN studies provided additional histological context for the pedunculopontine nucleus, ansa subthalamica and fibre-tract reconstruction^2,66,67^. These sources guide anatomical interpretation; they do not constitute an independent validation of every SWAN boundary or imply that every cited atlas mask was automatically transferred into the specimen.

The supplied segmentations are intended as an initial, revisable anatomical resource rather than definitive boundaries for every application. Any user can return to the source histology, edit an existing label or create a new one, and reapply the transformation chain to obtain corresponding labels in MRI or MNI space. Retaining editable geometry and the original images makes this process transparent and reproducible. This ability to review and extend the atlas is a central advantage of SWAN, particularly where tissue boundaries are uncertain or alternative anatomical definitions are relevant.

### Previous publications using the dataset

The present dataset has already supported several publications that correlated anatomical information provided by histological stacks with clinical findings. Examples include post-operative localizations of DBS electrodes within planned DBS targets in the subthalamic nucleus^68^, microdialysis of the globus pallidus internus^69,70^ and use of the spatial 3D atlas locations to propose a single trajectory to target both thalami and caudal zona incerta to treat essential tremor^71^. Display of thalamic nuclei depicting structural effect of the isodose curves for gamma ray radiosurgery for complex pain syndromes were also possible using normalized segmentations derived from the present resource^72^. Populational multicenter studies evaluating basal forebrain atrophy in Alzheimer’s disease^73^, primary progressive aphasia^74^ and predicting cognitive decline in *de novo* Parkinson’s disease^75^ were performed using basal forebrain histological masks obtained from the atlas. The pedunculopontine nucleus (PPN) mask published by our group^66^ allowed to estimate clinical improvements, evaluating cholinergic deficiency in cervical dystonia^76^ and to predict postural and gait symptoms in Parkinson’s Disease by extraction of neuroimaging metrics in 147 participants of the Parkinson’s Progression Markers Initiative^77^. Further work involved detailed correlation of lateral hypothalamic nuclei and local field potentials in hunger and satiety states in humans ^78^, safety and outcome of lateral hypothalamic DBS for obesity^79^, DBS modulation of fibers in cluster headache^80^, the illustration of anatomical structures involved in aggression^81^ and DBS in Tourette’s syndrome^82^. Finally, purely anatomical publications illustrated Edinger comb system^83^ and described a neglected fiber tract^2^ that may have clinical importance in DBS-induced psychiatric symptoms^84^.

Given these initial successes, we believe that making the atlas openly available together with software tools to enhance usability as a community resource will promote further translational work of the same kind by others. Contributions from other researchers to further improve the dataset, such as in form of additional segmentations of structures of interest are welcome and may be added to the repository. In figure 8, we show examples of brainstem regions included in the dataset, based on which brainstem nuclei and fiber tracts may be clearly appreciated. The fiber tracts segmented in the histological masks can be visualized using Histological Mesh Tractography (HMT), described elsewhere^67^. Such reconstructions are shown in Figure 9 (d).

**Figure 8.**
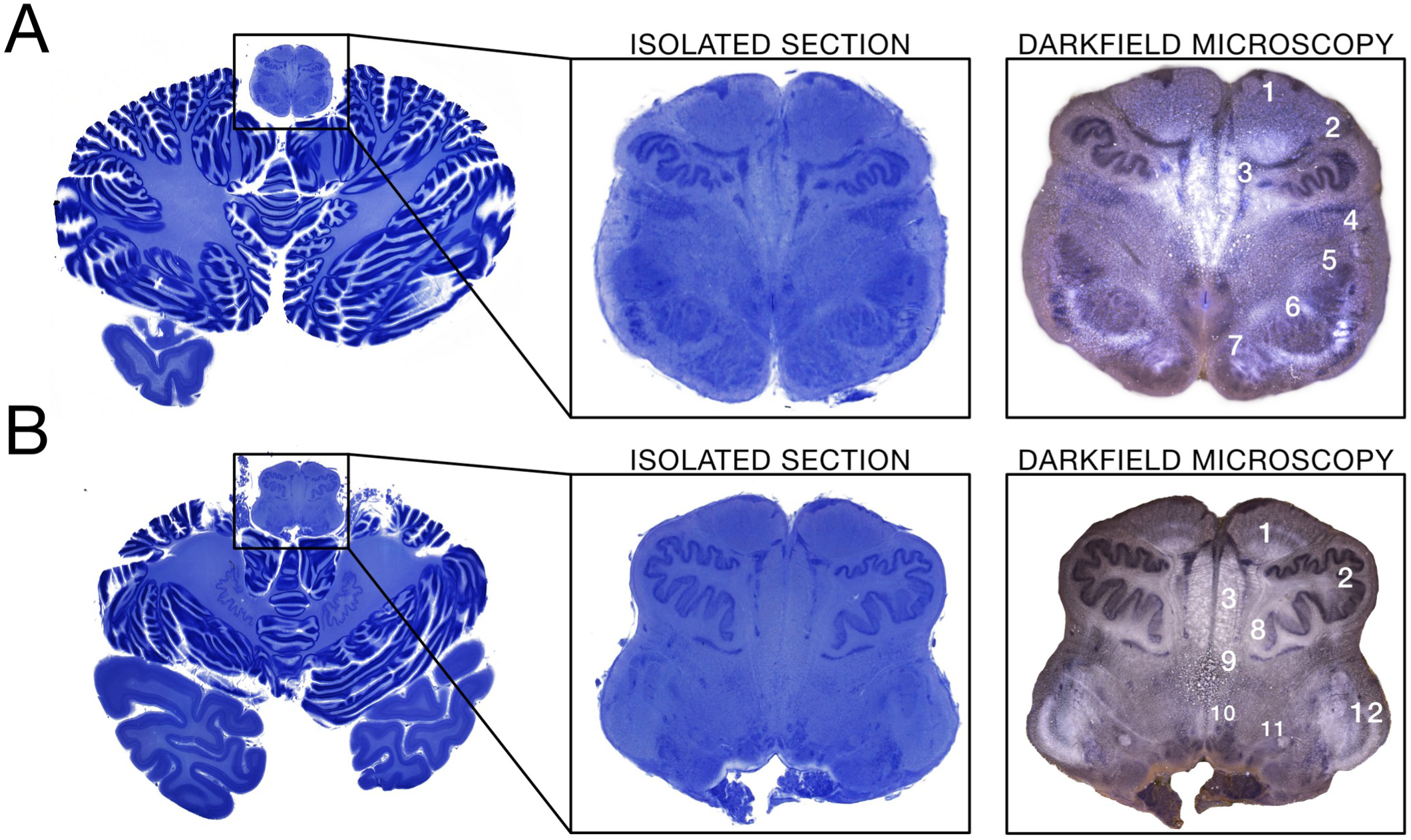
Bright field sections through the medulla oblongata and the cerebellum at a more (A) inferior and (B) superior level. Left panel: Magnified isolated sections. Right panel: Corresponding darkfield microscopy images showcasing even finer cytoarchitectonic details than seen in conventional microscopy. 1: Pyramids, 2: Inferior Olive, 3: Medial lemniscus, 4: Arcuate fibers, 5: Spinal nucleus of the trigeminus, 6: Nucleus cuneatus, 7: Nucleus gracilis, 8: Cerebello-olivary fibers, 9: Medial longitudinal fasciculus, 10: Hypoglossal nucleus, 11: Vagus nucleus, 12: Restiform body.

**Figure 9.**
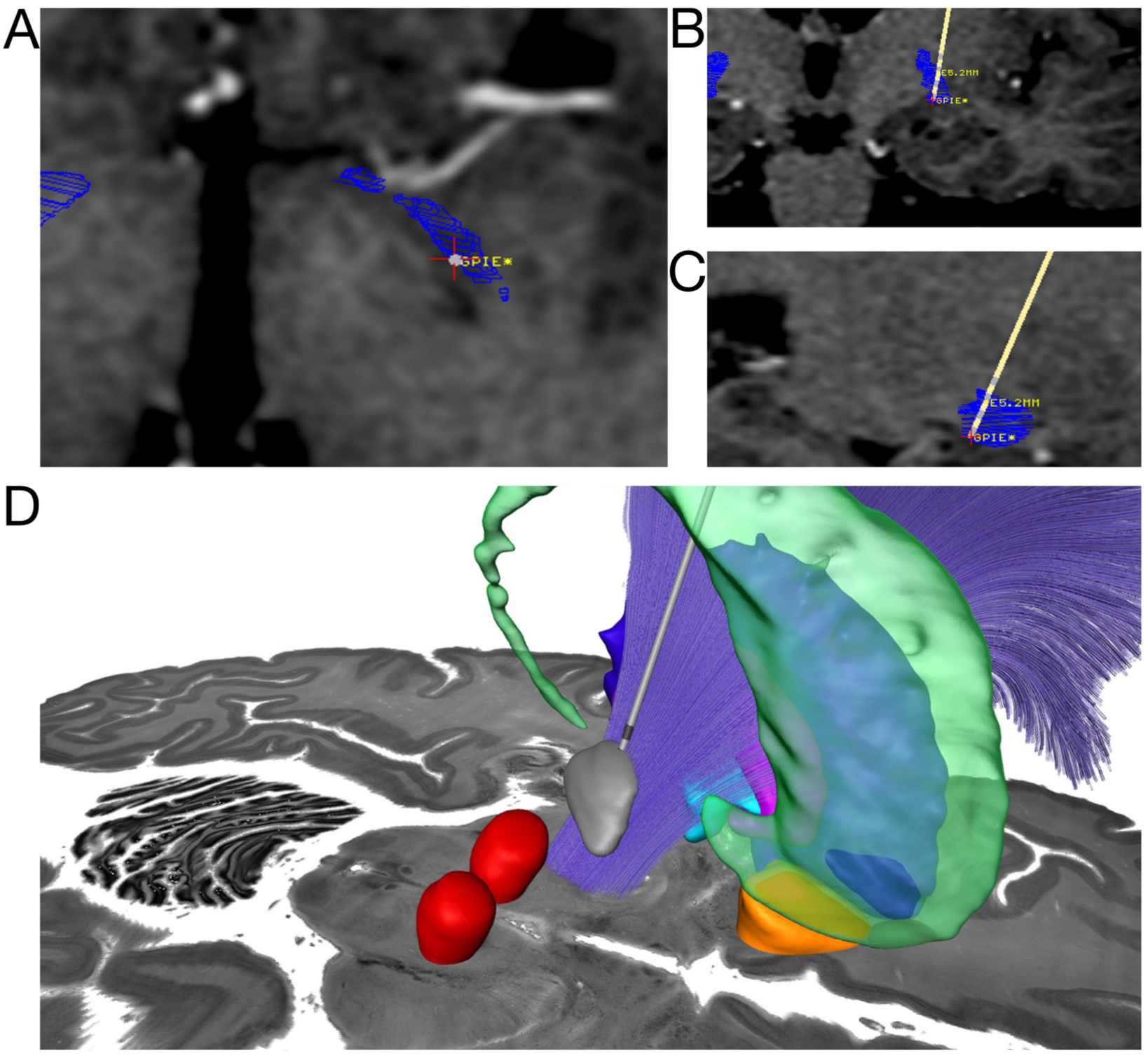
Stereotactic planning of GPi DBS for dystonia in MNPS using as confirmation tool the histological outlines defined in the SWAN atlas (A) axial plane (B) coronal plane and (C) sagittal plane (D) Post-operative reconstruction of a STN DBS merged into the SWAN atlas and histological background.

### Integration with clinical and research software

Our main goal while developing the SWAN atlas was to apply it to DBS planning and evaluation of postoperative DBS electrodes. In this context, the atlas integration in software solutions is essential.

For stereotactic planning, we have integrated the atlas into the Mevis Neurosurgery Planning System - MNPS (www.mevis.com.br). This system is suited for planning of functional stereotactic procedures, biopsies, stereotactic brachytherapy and radiosurgery. It allows multimodal imaging fusion (CT, MRI, stereotactic angiography and PET). MNPS is certified by the Brazilian National Agency of Health Surveillance (ANVISA), under license number 80195520021, for stereotactic planning of neurological surgeries in humans in Brazil. Although ready to use in a beta version, the SWAN atlas is currently not available in the commercial versions of MNPS.

The main advantage of using the SWAN for stereotactic planning either in MNPS or other software is that, since histological labels are registered to its own MRI, the fusion of the atlas to the patient is performed by coregistering the atlas MRI to the patient’s MRI. This is done by non-linear mutual information algorithms that are robust and less prone to errors than the previously used alternative of simply applying affine transforms to landmark points such as anterior commissure, posterior commissure or fastigial point (Figure 9 A, B and C).

Regarding postoperative analysis, the SWAN was further integrated into Lead-DBS (www.lead-dbs.org/^56^), which is a commonly used open source toolbox facilitating DBS electrode reconstructions and computer simulations based on postoperative MRI and CT imaging. Here, it serves as an atlas to visualize electrode reconstructions relative to anatomical structures. Lead-DBS is a research software not intended for clinical use but can be informative to investigate relationships between electrode placement and clinical improvements in a research setting. By use of the atlas, DBS electrodes may be localized to specific anatomical structures after surgery. Local corrections for better atlas fitting inside Lead-DBS environment can be achieved with WarpDrive^59^ tool.

Besides both software tools described above, we are developing 4 SWAN-dedicated open 3D Slicer modules to facilitate users to interact with the atlas capabilities. These tools are:

**SwanWarp:** is an interactive registration module for automatic and manual refinement of volumetric alignments between SWAN histology, patient MRI and stereotactic reference spaces. It was originally developed for the histology to MRI registration and to help the atlas MRI to MNI space normalization. It uses paired anatomical landmarks and stabilizing anchor points to generate localized nonlinear corrections while preserving the overall geometry of the registered volume. This module, besides its original use, can improve the registration of the SWAN atlas to patient MRI, despite non-linear complex differences between the atlas brain and the patient’s brain.

**SwanViewer:** provides an integrated interface for visualizing SWAN histology, blockface, MRI, MNI-space data and derived anatomical labels. The module is designed to support multimodal inspection, atlas–patient comparison and navigation across corresponding anatomical spaces without requiring users to manage the underlying transform chain manually.

**SwanLabel:** provides a dedicated environment for creating, editing and reviewing anatomical segmentations in high-resolution SWAN blockface/histology space. Segmentations can be stored with versioned metadata and propagated through the official transform chain to subject MRI and MNI spaces for downstream visualization and analysis. Segmentations in patient or MNI space can also be visualized in high-resolution and color histological space through application of the inverse transforms.

A fourth module is planned, but still not operational, the **SwanDBSPlanner**. It is designed for stereotactic planning using the patient’s MRI together with SWAN-derived anatomical overlays. The module will support AC–PC definition, target and trajectory planning, electrode-model placement and visualization of selected atlas structures in patient or stereotactic space. By using it together with the registration modules, an accurate fusion of the histological labels into patient space can enhance the understanding of the individual anatomy.

The distribution separates a lightweight core package from optional high-resolution asset packs. The core package contains code, manifests, transform definitions, low-resolution volumes, segmentation sources and final labels. Optional downloads contain high-resolution blockface-space and final-MNI-space intensity volumes. This keeps routine label use feasible on low-RAM computers while preserving high-resolution SWAN volumes as the central medical and scientific asset.

The public package is designed for a simple clinical-research workflow in 3D Slicer or Lead-DBS: load the relevant MRI, blockface or MNI background; load existing labels or author new segmentations in high-resolution blockface space; run the packaged forward or inverse label scripts; and inspect the result without manually managing the complete chain of intermediate transforms.

### Education

Besides the clinical and surgical applications of a 3D histological map warped into a stereotactic common space, we foresee further applications in education. To facilitate this process, we introduce a prototypical software tool called Neuroanatomy VR with this publication. The tool may be used to better understand the 3D relationships between brain structures. It is possible to navigate on both Android or Apple platforms, or to use it with simple VR glasses such as the Google Cardboard. Supplementary video 1 and figure 10 document the ongoing development of this educational software project.

**Figure 10.**
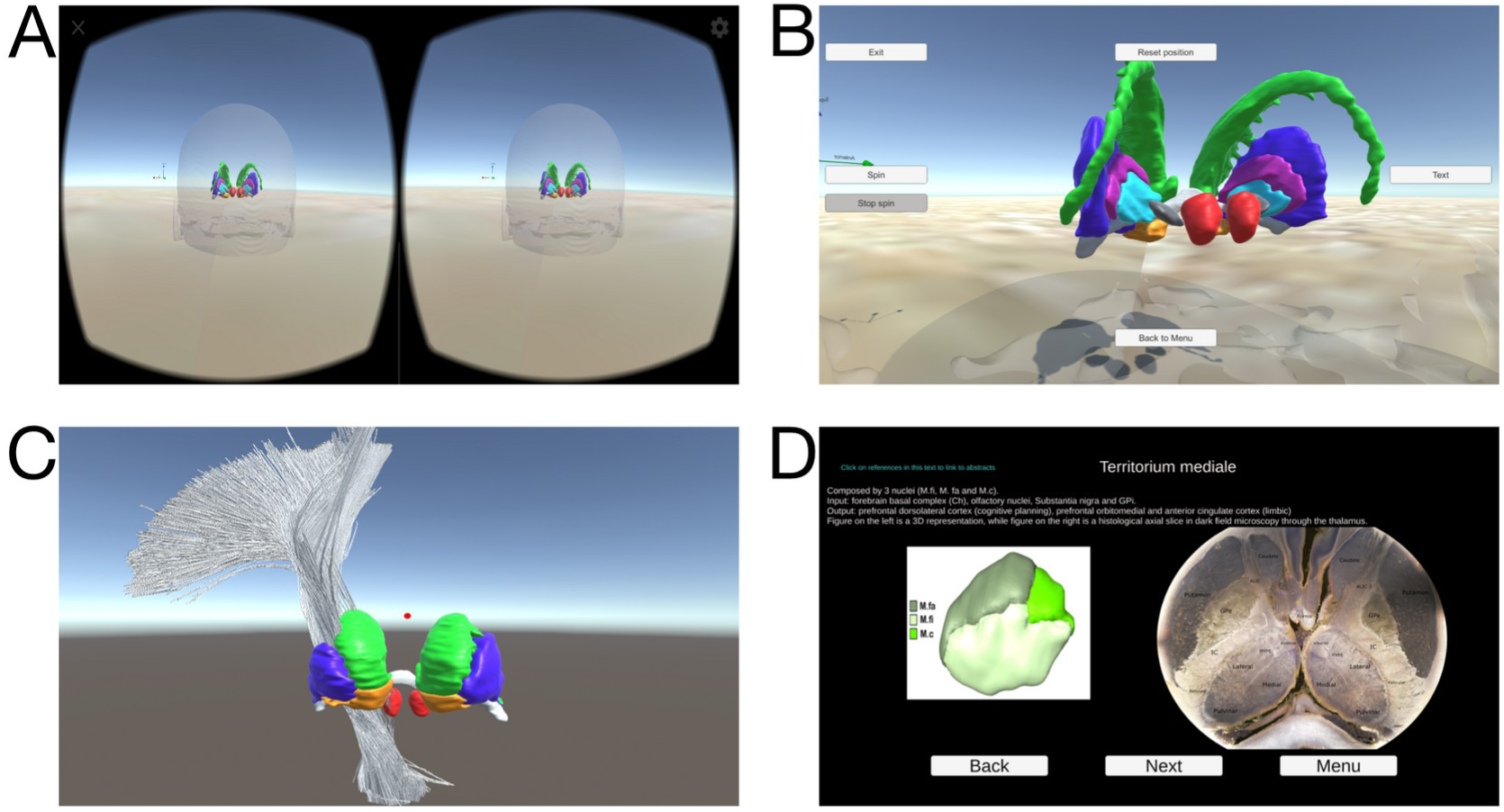
NeuroanatomyVR is a software project, aiming to use 3D histological segmentations as a resource to understand subcortical anatomy. It is possible to navigate through the structures and see histological slices from which the 3D forms are derived. Texts and references are directly linked to Pubmed abstracts.

## Discussion

Atlasing the human brain has been an endeavor of neuroscience since centuries. Histological staining of slices is considered the ground truth method to delineate and define micro- and mesoscopic structures and throughout history, multiple atlases have been created. However, not many datasets exist for which an entire human brain has been sectioned in standardized (i.e. continuous and equispaced) fashion, with a key dataset to mention the BigBrain project led by the center in Jülich^24^. With this report, we describe a whole-brain dataset with systematic slicing of comparably thick sections. The choice for thick sections was a deliberate one, since it led to far fewer nonlinear distortions of slices and hence fewer issues in 2D and 3D registrations, during downstream registration. The registered volumes of various MRI sequences, including DTI, with different histological modes (blockface, darkfield and lightfield) generates a multimodal contrast resource, that could potentially be used to improve automatic segmentation algorithms (Figure 11).

**Figure 11.**
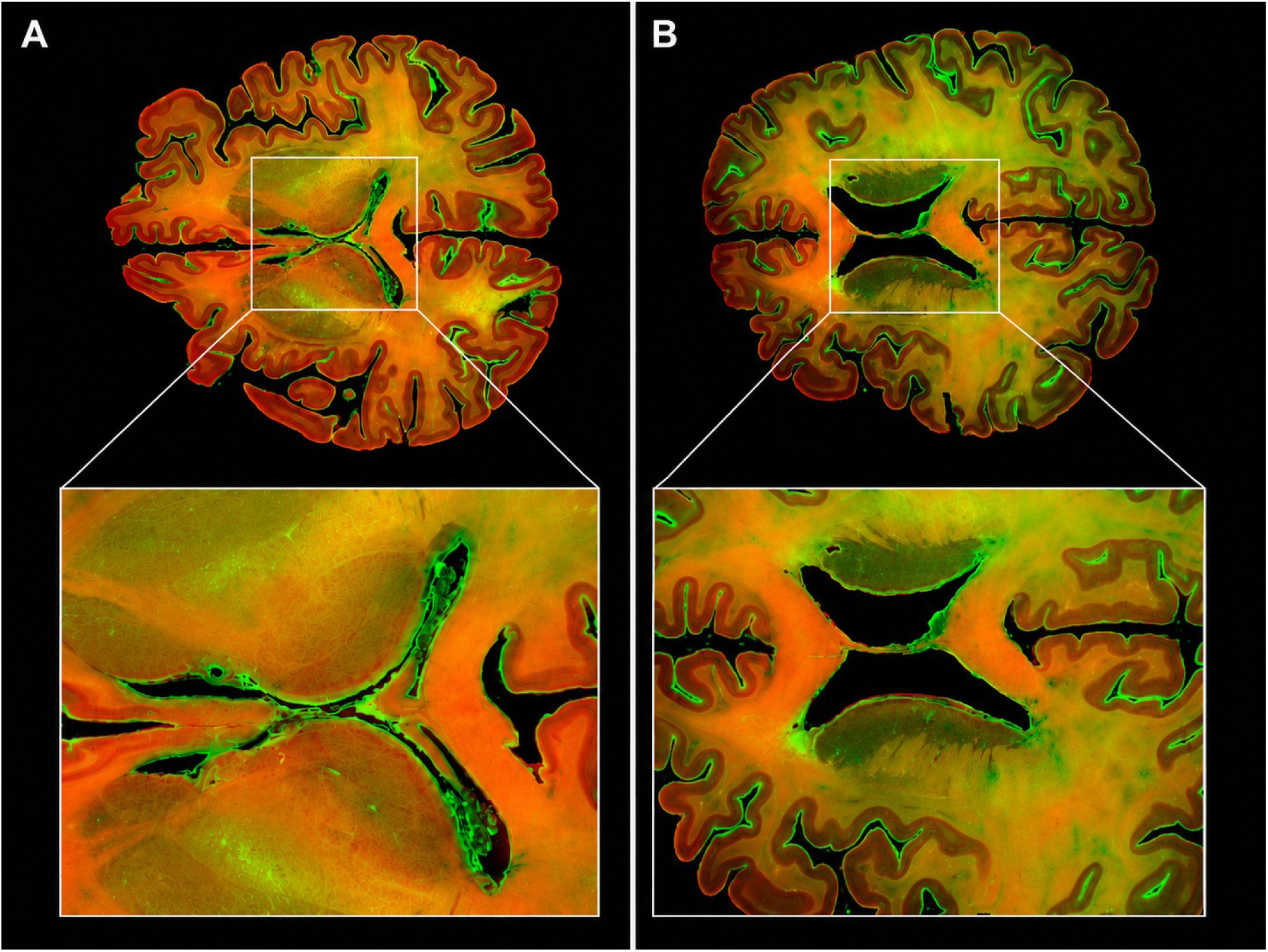
The superposition of blockface images (red) and Nissl stained histological slices (green) can enhance the properties of each modality, showing an interesting contrast between white matter and gray matter. In (A) it is possible to clearly visualize the contrast in the anterior thalamic radiation, crossing the head of caudate nucleus, connecting to the medial thalamus through its anterior region. In (B), the forceps major and forceps minor are enhanced by the multimodal histological imaging

Our dataset has been created by a team of neurosurgeons and anatomists, which continuously kept the goal of creating a clinically useful resource in mind. Hence, meticulous detail has been spent on delineating deep structures involved in brain surgery, as well as accompanying darkfield photography of subcortical structures relevant for clinical interventions. Before sectioning, the brain was scanned *in-situ*, making it straight-forward to register segmentations to patient data. Moreover, we provide a software framework that will allow translation of segmentations from histology to standard stereotactic space and back. Using this tool, for instance, one could apply findings from neuroimaging studies (e.g., functional MRI results, brain-lesion sites or meta-analytic results) and translate them back into the space of histological sections to help identify the exact location of such findings relative to histological landmarks.

Together, the atlas and accompanying software tools provide a unique dataset that may be helpful for both clinical, translational, and basic science endeavors. Our prior results show utility of the tool and atlas in functional stereotactic surgery^2,48,66–74,76–80,84^.

BigBrain reconstructed 7,404 histological sections at 20 µm isotropic resolution^24^. SWAN uses 430 µm-thick sections with 25 µm in-plane image sampling, giving it substantially coarser through-plane sampling. The use of thick sections was intended to reduce section-level deformation and facilitate registration to blockface images and the specimen MRI^48^. The two resources therefore provide complementary anatomical information. SWAN emphasizes the correspondence among serial thick-section histology, blockface imaging and in situ MRI, together with editable subcortical labels and reversible transformations into stereotactic space. These features support anatomical inspection and atlas adaptation, but do not imply spatial resolution equivalent to BigBrain. Subcortical structures were delineated through close collaboration between neurosurgeons and neuroanatomists.

As limitations, SWAN lacks ultrahigh *in-plane* resolution scans (although possible to process them in a future release), and the dataset does not include specialized fibre staining or immunostaining. Furthermore, darkfield scans have been acquired with the aim to cover most relevant brain regions but for technical reasons had to be acquired with a comparatively smaller field of view. This limitation also can be overcome in a future release, since the original data is preserved. However, meticulous registrations of the slices to each other and to the light-field (whole-brain) scans made it possible to still register darkfield slices into a stack/volume space. The dataset also comprises a single brain. While typical for histological atlases, this limits its applicability to populations with pathology and to very young or old populations. This limitation can be minimized by using the non-linear deformation tools developed for the atlas fitting using its own MRI (SwanWarp).

## Conclusions

We describe a whole-brain histological deformable atlas dataset intended for open academic use. The dataset comprises dark- and light field microscopy data, an MRI of the brain scanned *in situ* before brain extraction and slicing, as well as tools to translate regions of interest from histological sections to standard stereotactic space. The dataset has already supported numerous publications in the field of stereotactic surgery and beyond. With its planned open release, we believe that it may enable a larger number of future research studies.

## Data availability

At the time of this preprint, the SWAN data packages have not yet been deposited in public repositories and no dataset DOIs have been assigned. The project website (https://www.swanatlas.com) and its Open Data page (https://www.swanatlas.com/open-data/) describe the resource and will provide links to the staged public release. Zenodo (https://zenodo.org/) is the planned archival repository for the following packages: (i) a lightweight SwanAtlasPipeline core, including metadata, transform manifests and definitions, quality-control reports, preview volumes, editable segmentation sources and derived labels; (ii) high-resolution blockface-space blockface, brightfield grayscale/RGB and darkfield volumes; (iii) high-resolution final-MNI-space volumes; (iv) the native specimen MRI, DTI and redistributable DSI Studio derivatives; and (v) a Lead-DBS atlas package containing final-MNI labels. MRI/DTI derivatives may be included within the core package or deposited separately according to size and redistribution permissions. Separate records are planned for the high-resolution packages. Repository-specific URLs and persistent identifiers will be added when these deposits are available.

## Code availability

Source code for SwanAtlasPipeline and its 3D Slicer modules is planned for a public GitHub repository (https://github.com/), with versioned releases archived in Zenodo. The planned Lead-DBS runtime package will include portable transform manifests, official forward/inverse transforms, lightweight reference images, MATLAB and shell/Python wrappers, test examples and documentation. Neuroanatomy VR will be distributed separately as an educational companion, with Android/Mac builds and cleaned Unity source archived in a dedicated Zenodo record. No SWAN-specific GitHub URL, software deposit or software DOI has yet been assigned. Release links, licences, version information and checksums will be indexed on the SWAN website as they become available. SwanDBSPlanner remains a planned module and is not an operational release.

## Supporting information

Table 1 - SWAN anatomical structures and subdivisions

Supplementary Video 1 - NeuroanatomyVR demonstration

## Acknowledgements

We thank the donor’s family and all the team of the Brazilian Brain Bank of the Aging Brain Study Group for enabling this work.

## Author contributions

E.J.L.A. and H.H. conceived the SWAN atlas resource. E.J.L.A., A.T.D.L.A., E.A.J., L.T.G., E.T.F. and H.H. contributed to specimen acquisition, MRI, histology, segmentation strategy or clinical framing. H.F., S.O. and A.H. contributed to stereotactic registration, Lead-DBS/WarpDrive integration and spatial normalization strategy. A.A.B. contributed to MNPS/software integration. E.J.L.A. coordinated the current pipeline reconstruction and manuscript preparation.

## References

1. Edlow, B. L. et al. 7 Tesla MRI of the ex vivo human brain at 100 micron resolution. Sci. Data 6, 244 (2019).

2. Alho, E. J. L. et al. The Ansa Subthalamica: A Neglected Fiber Tract. Mov. Disord. Off. J. Mov. Disord. Soc. 35, 75–80 (2020).

3. Horn, A. et al. Lead-DBS v2: Towards a comprehensive pipeline for deep brain stimulation imaging. NeuroImage 184, 293–316 (2019).

4. Spiegel EA, Wycis HT, & Freed H. Stereoencephalotomy: Thalamotomy and related procedures. J. Am. Med. Assoc. 148, 446–451 (1952).

5. Talairach, J. Atlas d’anatomie stéréotaxique; repérage radiologique indirect des noyaux gris centraux des régions mésencéphalo-sous-optique et hypothalamique de l’homme,. (Masson, Paris, 1957).

6. Schaltenbrand, G. & Bailey, P. Introduction to Stereotaxis with an Atlas of the Human Brain, Vol. I: Text; Vol. II: Plate 1-57, Vol. III: Plate 58-76. (Georg Thieme, Grune & Stratton, 1959).

7. Schaltenbrand, G., Hassler, R. & Wahren, W. Atlas for Stereotaxy of the Human Brain: With an Accompanying Guide. (Thieme, Stuttgart, 1977).

8. Andrew, J. & Watkins, E. S. A Stereotaxic Atlas of the Human Thalamus and Adjacent Structures: A Variability Study. (Williams & Wilkins, Baltimore, 1969).

9. Van Buren, J. M. & Borke, R. C. Variations and Connections of the Human Thalamus. (Springer-Verlag, Berlin; New York, 1972).

10. Emmers, R. & Tasker, R. The Human Somesthetic Thalamus : With Maps for Physiological Target Localization during Stereotactic Neurosurgery. (Raven Press, New York, 1975).

11. Afshar, F., Watkins, E. S. & Yap, J. C. Stereotaxic Atlas of the Human Brainstem and Cerebellar Nuclei : A Variability Study. (Raven Press, New York, 1978).

12. Yoshida, M. Creation of a three-dimensional atlas by interpolation from Schaltenbrand-Bailey’s atlas. Appl. Neurophysiol. 50, 45–48 (1987).

13. Talairach, J. & Tournoux, P. Co-Planar Stereotaxic Atlas of the Human Brain: 3-Dimensional Proportional System: An Approach to Medical Cerebral Imaging. (Thieme, Stuttgart [etc.], 1988).

14. Ono, M., Kubik, S. & Abernathey, C. D. Atlas of the Cerebral Sulci. (G. Thieme Verlag ; Thieme Medical Publishers, Stuttgart; New York, 1990).

15. Morel, A. Stereotactic Atlas of the Human Thalamus and Basal Ganglia. (Informa Healthcare, New York, 2007).

16. Morel, A., Magnin, M. & Jeanmonod, D. Multiarchitectonic and stereotactic atlas of the human thalamus. J. Comp. Neurol. 387, 588–630 (1997).

17. Nowinski, W. L. & Belov, D. The Cerefy Neuroradiology Atlas: a Talairach-Tournoux atlas-based tool for analysis of neuroimages available over the internet. NeuroImage 20, 50–57 (2003).

18. Nowinski, W. L., Belov, D., Thirunavuukarasuu, A. & Benabid, A. L. A probabilistic functional atlas of the VIM nucleus constructed from pre-, intra- and postoperative electrophysiological and neuroimaging data acquired during the surgical treatment of Parkinson’s disease patients. Stereotact. Funct. Neurosurg. 83, 190–196 (2005).

19. Chakravarty, M. M., Bertrand, G., Hodge, C. P., Sadikot, A. F. & Collins, D. L. The creation of a brain atlas for image guided neurosurgery using serial histological data. NeuroImage 30, 359–376 (2006).

20. Yelnik, J. et al. A three-dimensional, histological and deformable atlas of the human basal ganglia. I. Atlas construction based on immunohistochemical and MRI data. NeuroImage 34, 618–638 (2007).

21. Mai, J. K., Paxinos, G. & Voss, T. Atlas of the Human Brain. (Academic Press, New York; London, 2008).

22. Zaborszky, L. et al. Stereotaxic probabilistic maps of the magnocellular cell groups in human basal forebrain. NeuroImage 42, 1127–1141 (2008).

23. Krauth, A. et al. A mean three-dimensional atlas of the human thalamus: generation from multiple histological data. NeuroImage 49, 2053–2062 (2010).

24. Amunts, K. et al. BigBrain: an ultrahigh-resolution 3D human brain model. Science 340, 1472–1475 (2013).

25. König, J. F. R. & Klippel, R. A. The Rat Brain: A Stereotaxic Atlas of the Forebrain and Lower Parts of the Brain Stem. (Williams and Wilkins, 1963).

26. Paxinos, G. & Watson, C. The Rat Brain in Stereotaxic Coordinates. (Academic Press, 1986).

27. Franklin, K. & Paxinos, G. The Mouse Brain in Stereotaxic Coordinates. (Academic Press, 2008).

28. Kroon, J. P. & Riley, A. L. A microcomputer-based system for stereotaxic coordinates in the rat brain. Physiol. Behav. 38, 593–596 (1986).

29. Toga, A. W., Samaie, M. & Payne, B. A. Digital rat brain: a computerized atlas. Brain Res. Bull. 22, 323–333 (1989).

30. Schwarz, A. J. et al. A stereotaxic MRI template set for the rat brain with tissue class distribution maps and co-registered anatomical atlas: application to pharmacological MRI. NeuroImage 32, 538–550 (2006).

31. Kumazawa-Manita, N., Katayama, M., Hashikawa, T. & Iriki, A. Three-dimensional reconstruction of brain structures of the rodent Octodon degus: a brain atlas constructed by combining histological and magnetic resonance images. Exp. Brain Res. 231, 65–74 (2013).

32. Shanta, T. R., Manocha, S. L. & Bourne, G. H. A Stereotaxic Atlas of the Java Monkey Brain (Macaca irus). 2b–23 (1968) doi:10.1159/000390681.

33. François, C., Yelnik, J. & Percheron, G. A stereotaxic atlas of the basal ganglia in macaques. Brain Res. Bull. 41, 151–158 (1996).

34. Martin, R. F. & Bowden, D. M. A stereotaxic template atlas of the macaque brain for digital imaging and quantitative neuroanatomy. NeuroImage 4, 119–150 (1996).

35. Ilinsky, I. A., Kultas-Ilinsky, K. & Knosp, B. *Stereotactic Atlas of the Macaca Mulatta Thalamus and Adjacent Basal Ganglia Nuclei*. (Springer US, Boston, MA, 2002). doi:10.1007/978-1-4615-0625-6.

36. Saleem, K. S. & Logothetis, N. K. A Combined MRI and Histology Atlas of the Rhesus Monkey Brain in Stereotaxic Coordinates. (Academic Press, 2012).

37. Lanciego, J. L. & Vázquez, A. The basal ganglia and thalamus of the long-tailed macaque in stereotaxic coordinates. A template atlas based on coronal, sagittal and horizontal brain sections. Brain Struct. Funct. 217, 613–666 (2012).

38. Dauguet, J. et al. Three-dimensional reconstruction of stained histological slices and 3D non-linear registration with in-vivo MRI for whole baboon brain. J. Neurosci. Methods 164, 191–204 (2007).

39. Toga, A. W., Thompson, P. M., Mori, S., Amunts, K. & Zilles, K. Towards multimodal atlases of the human brain. Nat. Rev. Neurosci. 7, 952–966 (2006).

40. Friedrich, H. et al. A precise atlas of the human subcortex. bioRxiv [preprint] (2026). 10.64898/2026.02.13.705755.

41. Mai, J. K. & Majtanik, M. Toward a Common Terminology for the Thalamus. Front. Neuroanat. 12, (2019).

42. Niemann, K. & van Nieuwenhofen, I. One Atlas-Three Anatomies: Relationships of the Schaltenbrand and Wahren Microscopic Data. Acta Neurochir. (Wien) 141, 1025–1038 (1999).

43. Niemann, K., Naujokat, C., Pohl, G., Wollner, C. & von Keyserlingk, D. Verification of the Schaltenbrand and Wahren stereotactic atlas. Acta Neurochir. (Wien) 129, 72–81 (1994).

44. Nowinski, W. L., Liu, J. & Thirunavuukarasuu, A. Quantification and visualization of the three-dimensional inconsistency of the subthalamic nucleus in the Schaltenbrand-Wahren brain atlas. Stereotact. Funct. Neurosurg. 84, 46–55 (2006).

45. Nowinski, W. L. et al. Multiple brain atlas database and atlas-based neuroimaging system. Comput. Aided Surg. Off. J. Int. Soc. Comput. Aided Surg. 2, 42–66 (1997).

46. Heinsen, H. & Heinsen, Y. L. Serial Thick, Frozen, Gallocyanin Stained Sections of Human Central Nervous System. J. Histotechnol. 14, 167–173 (1991).

47. Heinsen, H., Arzberger, T. & Schmitz, C. Celloidin mounting (embedding without infiltration) - a new, simple and reliable method for producing serial sections of high thickness through complete human brains and its application to stereological and immunohistochemical investigations. J. Chem. Neuroanat. 20, 49–59 (2000).

48. Alho, E. J. L. et al. High thickness histological sections as alternative to study the three-dimensional microscopic human sub-cortical neuroanatomy. Brain Struct. Funct. 10.1007/s00429-017-1548-2 (2017) doi:10.1007/s00429-017-1548-2.

49. Grinberg, L. T. et al. Brain bank of the Brazilian aging brain study group - a milestone reached and more than 1,600 collected brains. Cell Tissue Bank. 8, 151 –162 (2007).

50. Alegro, M. et al. Multimodal Whole Brain Registration: MRI and High Resolution Histology. http://www.cv-foundation.org/openaccess/content_cvpr_2016_workshops/w15/papers/Alegro_Multimodal_Whole_Brain_CVPR_2016_paper.pdf (2016).

51. Yang, J., Lu, W. & Waibel, A. Skin-color modeling and adaptation. in Computer Vision — ACCV’98 (eds Chin, R. & Pong, T.-C.) 687–694 (Springer, Berlin, Heidelberg, 1997). doi:10.1007/3-540-63931-4_278.

52. ANTs by stnava. http://stnava.github.io/ANTs/.

53. Mattes, D., Haynor, D. R., Vesselle, H., Lewellen, T. K. & Eubank, W. PET-CT image registration in the chest using free-form deformations. IEEE Trans. Med. Imaging 22, 120–128 (2003).

54. Sled, J. G., Zijdenbos, A. P. & Evans, A. C. A nonparametric method for automatic correction of intensity nonuniformity in MRI data. IEEE Trans. Med. Imaging 17, 87–97 (1998).

55. Avants, B. B., Epstein, C. L., Grossman, M. & Gee, J. C. Symmetric diffeomorphic image registration with cross-correlation: evaluating automated labeling of elderly and neurodegenerative brain. Med. Image Anal. 12, 26–41 (2008).

56. Horn, A. & Kühn, A. A. Lead-DBS: A toolbox for deep brain stimulation electrode localizations and visualizations. NeuroImage 107, 127–135 (2015).

57. Ewert, S. et al. Optimization and comparative evaluation of nonlinear deformation algorithms for atlas-based segmentation of DBS target nuclei. NeuroImage 184, 586–598 (2019).

58. Vogel, D. et al. Anatomical brain structures normalization for deep brain stimulation in movement disorders. NeuroImage Clin. 27, 102271 (2020).

59. Oxenford, S. et al. WarpDrive: Improving spatial normalization using manual refinements. Med. Image Anal. 91, 103041 (2024).

60. Amunts, K., Mohlberg, H., Bludau, S. & Zilles, K. Julich-Brain: A 3D probabilistic atlas of the human brain’s cytoarchitecture. Science 369, 988–992 (2020). 10.1126/science.abb4588.

61. Amunts, K. et al. Julich-Brain Atlas, cytoarchitectonic maps (v3.1). EBRAINS [dataset], version 3.1 (2024). 10.25493/KNSN-XB4.

62. Amunts, K. et al. Cytoarchitectonic mapping of the human amygdala, hippocampal region and entorhinal cortex: intersubject variability and probability maps. Anat. Embryol. 210, 343–352 (2005). 10.1007/s00429-005-0025-5.

63. Kedo, O. et al. Receptor-driven, multimodal mapping of the human amygdala. Brain Struct. Funct. 223, 1637–1666 (2018). 10.1007/s00429-017-1577-x.

64. Brandstetter, A. et al. Organization, fine structure, and stereotaxic maps of the human Bed nucleus of the Stria terminalis. Imaging Neurosci. 4, IMAG.a.1260 (2026). 10.1162/imag.a.1260.

65. Tellmann, S. et al. Cytoarchitectonic mapping of the human brain cerebellar nuclei in stereotaxic space and delineation of their co-activation patterns. Front. Neuroanat. 9, 54 (2015). 10.3389/fnana.2015.00054.

66. Alho, A. T. D. L. et al. Magnetic resonance diffusion tensor imaging for the pedunculopontine nucleus: proof of concept and histological correlation. Brain Struct. Funct. 10.1007/s00429-016-1356-0 (2017) doi:10.1007/s00429-016-1356-0.

67. Alho, E. J. L., Fonoff, E. T., Di Lorenzo Alho, A. T., Nagy, J. & Heinsen, H. Use of computational fluid dynamics for 3D fiber tract visualization on human high-thickness histological slices: histological mesh tractography. Brain Struct. Funct. 10.1007/s00429-020-02187-3 (2021) doi:10.1007/s00429-020-02187-3.

68. Fonoff, E. T., Campos, W. K., Mandel, M., Alho, E. J. L. & Teixeira, M. J. Bilateral subthalamic nucleus stimulation for generalized dystonia after bilateral pallidotomy. Mov. Disord. Off. J. Mov. Disord. Soc. 27, 1559–1563 (2012).

69. Lopez, W. O. C., et al. Optimizing microdialysis for deep brain stimulation. Front. Biosci. Elite Ed. 8, 299–310 (2016).

70. Martinez, R. C. R. et al. Intraoperative dopamine release during globus pallidus internus stimulation in Parkinson’s disease. Mov. Disord. Off. J. Mov. Disord. Soc. 10.1002/mds.25691 (2013) doi:10.1002/mds.25691.

71. Dos Santos Ghilardi, M. G., et al. Double-target DBS for essential tremor: 8-contact lead for cZI and Vim aligned in the same trajectory. Neurology 90, 476–478 (2018).

72. Lovo, E. E. et al. Stereotactic Gamma Ray Radiosurgery to the Centromedian and Parafascicular Complex of the Thalamus for Trigeminal Neuralgia and Other Complex Pain Syndromes. Cureus 11, e6421 (2019).

73. Kilimann, I. et al. Subregional basal forebrain atrophy in Alzheimer’s disease: a multicenter study. J. Alzheimers Dis. JAD 40, 687–700 (2014).

74. Teipel, S. J. et al. Brain atrophy in primary progressive aphasia involves the cholinergic basal forebrain and Ayala’s nucleus. Psychiatry Res. 221, 187–194 (2014).

75. Ray, N. J. et al. In vivo cholinergic basal forebrain atrophy predicts cognitive decline in de novo Parkinson’s disease. Brain J. Neurol. 141, 165–176 (2018).

76. Mente, K. et al. Pedunculopontine Nucleus Cholinergic Deficiency in Cervical Dystonia. Mov. Disord. Off. J. Mov. Disord. Soc. 33, 827–834 (2018).

77. Craig, C. E. et al. Pedunculopontine Nucleus Microstructure Predicts Postural and Gait Symptoms in Parkinson’s Disease. Mov. Disord. **n/a**, (2020).

78. Talakoub, O. et al. Lateral hypothalamic activity indicates hunger and satiety states in humans. Ann. Clin. Transl. Neurol. 4, 897–901 (2017).

79. Franco, R. R. et al. Assessment of Safety and Outcome of Lateral Hypothalamic Deep Brain Stimulation for Obesity in a Small Series of Patients With Prader-Willi Syndrome. *JAMA Netw*. Open 1, (2018).

80. Dantas, S. A. F. et al. Deep brain stimulation modulates hypothalamic-brainstem fibers in cluster headache: case report. J. Neurosurg. 1–4 (2019) doi:10.3171/2018.11.JNS181412.

81. Gouveia, F. V. et al. Amygdala and Hypothalamus: Historical Overview With Focus on Aggression. Neurosurgery 85, 11–30 (2019).

82. Casagrande, S. C. B., Cury, R. G., Alho, E. J. L. & Fonoff, E. T. Deep brain stimulation in Tourette’s syndrome: evidence to date. Neuropsychiatr. Dis. Treat. 15, 1061–1075 (2019).

83. Horn, A. et al. Teaching NeuroImages: In vivo visualization of Edinger comb and Wilson pencils. Neurology 92, e1663–e1664 (2019).

84. Gouveia, F. V. et al. The ansa subthalamica as a substrate for DBS-induced manic symptoms. Brain Stimulat. 10.1016/j.brs.2020.07.017 (2020) doi:10.1016/j.brs.2020.07.017.

