## Supplementary material for "The SWAN atlas: an open multimodal resource combining neuroimaging and high-resolution histology of the human brain in stereotactic space": Table 1 - SWAN anatomical structures and subdivisions

### SWAN anatomical structures and abbreviations

Supplementary Material for The SWAN atlas. Anatomical structures, regional complexes and pathways included in the SWAN structure list. Whole structures and their subdivisions are listed separately; abbreviations identify anatomical labels and do not imply a count of independent or non-overlapping regions.

| Anatomical structure | Abbreviation |
| --- | --- |
| **Basal ganglia and associated nuclei** | |
| Caudate nucleus | Cd |
| Putamen | Put |
| Nucleus accumbens | NAcc |
| Globus pallidus, internal segment | GPi |
| Globus pallidus, external segment | GPe |
| Subthalamic nucleus | STN |
| Claustrum | Cl |
| Bed nucleus of the stria terminalis, dorsal division | BNSTd |
| **Basal forebrain cholinergic fields** | |
| Medial septal nucleus | CH1 |
| Nucleus of the diagonal band of Broca, vertical limb | CH2 |
| Nucleus of the diagonal band of Broca, horizontal limb | CH3 |
| Nucleus basalis of Meynert | CH4 |
| **Amygdala and transition regions** | |
| Amygdala, whole structure | Amy |
| Central amygdaloid nucleus | CE |
| Lateral amygdaloid nucleus | LA |
| Basal amygdaloid nuclear region | BA |
| Accessory basal amygdaloid nucleus | AB |
| Medial and cortical amygdaloid nuclear group | ME |
| Corticoamygdaloid transition area | CAT |
| **Hypothalamus and mammillary region** | |
| Medial preoptic nucleus | MPO |
| Anterior hypothalamic area | AHA |
| Suprachiasmatic nucleus | SCN |
| Supraoptic nucleus | SON |

| Anatomical structure | Abbreviation |
|---|---|
| Paraventricular nucleus | PVN |
| Periventricular nucleus | Pe |
| Dorsomedial hypothalamic nucleus | DMH |
| Ventromedial hypothalamic nucleus | VMH |
| Arcuate nucleus | ARC |
| Lateral hypothalamic area | LH |
| Tuberomammillary nucleus | TMN |
| Posterior hypothalamic area | PH |
| Posterior hypothalamic nucleus | PHN |
| Mammillary bodies | MM |
| **Thalamus and metathalamus** | |
| Thalamus, whole structure | Thal |
| Reticular nucleus of the thalamus | Rt |
| Nucleus lateropolaris thalami | L.po |
| Nucleus anterior principalis | A.pr |
| Mediodorsal thalamic nucleus | MD |
| Nucleus centralis thalami | Ce |
| Nucleus parafascicularis | Pf |
| Nucleus dorsalis superficialis | D.sf |
| Nucleus limitans | Li |
| Nucleus dorso-oralis externus | D.o.e |
| Nucleus dorso-oralis internus | D.o.i |
| Nucleus dorso-intermedius externus | D.im.e |
| Nucleus dorso-intermedius internus | D.im.i |
| Nucleus dorso-caudalis | D.c |
| Nucleus zentrolateralis oralis | Z.o |
| Nucleus zentrolateralis intermedius | Z.im |
| Zentrointermedius externus / zentrolateralis externus | Z.im.e |
| Nucleus zentro-intermedius internus | Z.im.i |
| Nucleus zentro-caudalis externus | Z.c.e |
| Nucleus zentro-caudalis internus | Z.c.i |
| Nucleus ventro-oralis anterior | V.o.a |
| Nucleus ventro-oralis posterior | V.o.p |

| **Anatomical structure** | **Abbreviation** |
|---|---|
| Nucleus ventro-oralis internus | V.o.i |
| Nucleus ventro-oralis medialis | V.o.m |
| Nucleus ventro-intermedius externus | V.im.e |
| Nucleus ventro-intermedius internus | V.im.i |
| Nucleus ventro-caudalis externus | V.c.e |
| Nucleus ventro-caudalis internus | V.c.i |
| Nucleus ventro-caudalis anterior internus | V.c.a.i |
| Nucleus ventro-caudalis parvocellularis | V.c.pc |
| Nucleus ventro-caudalis parvocellularis externus | V.c.pc.e |
| Nucleus ventro-caudalis posterior externus | V.c.p.e |
| Nucleus ventro-caudalis portae | V.c.por |
| Nucleus pulvinaris orolateralis | Pu.o.l |
| Nucleus pulvinaris oromedialis | Pu.o.m |
| Nucleus pulvinaris oroventralis | Pu.o.v |
| Nucleus pulvinaris intergeniculatus | Pu.ig |
| Pulvinar mediale | Pu.m |
| Pulvinar laterale | Pu.l |
| Nucleus pulvinaris superficialis | Pu.sf |
| Medial geniculate body | G.m |
| **Deep cerebellar nuclei** | |
| Dentate nucleus | DN |
| Emboliform nucleus | EmN |
| Globose nucleus | GlN |
| Fastigial nucleus | FN |
| **Brainstem nuclei and regional complexes** | |
| Red nucleus | RN |
| Substantia nigra complex | SN |
| Ventral tegmental area complex | VTA |
| Superior colliculus | SC |
| Inferior colliculus | IC |
| Periaqueductal gray | PAG |
| Pedunculopontine / pedunculotegmental nucleus | PPN / PTg |
| Mesencephalic reticular region, including the cuneiform region | mRt |

| Anatomical structure | Abbreviation |
|---|---|
| Laterodorsal tegmental nucleus | LDTg |
| Locus coeruleus, noradrenergic group A6 | LC / A6 |
| Parabrachial complex | PBC |
| Pontine reticular nucleus, oral part | PnO |
| Dorsal raphe nucleus | DR |
| Median raphe nucleus | MnR |
| Inferior olivary complex | IO |
| Solitary nucleus complex | NTS |
| **Pallidal pathways and fields of Forel** | |
| Ansa lenticularis | AL |
| Ansa subthalamica | AS |
| Lenticular fasciculus and field H2 of Forel | FL / H2 |
| Thalamic fasciculus, field H1 of Forel | FT / H1 |
| Field H of Forel | H |
| Subthalamic fasciculus | SF |
| **Brainstem and cerebellar pathways** | |
| Dentatorubrothalamic tract, crossed component | cDRTT |
| Dentatorubrothalamic tract, nondecussating component | ndDRTT |
| Medial lemniscus | ML |
| Lateral lemniscus | LL |
| Dorsal longitudinal fasciculus | DLF |
| Medial longitudinal fasciculus | MLF |
| Central tegmental tract | CTT |
| Spinothalamic tract | STT |
| Corticospinal tract, pyramidal pathway | CST |
| Superior cerebellar peduncle | SCP |
| Middle cerebellar peduncle | MCP |
| Inferior cerebellar peduncle | ICP |
| **Limbic pathways and other fiber bundles** | |
| Fornix | Fx |
| Mammillothalamic tract | MTT |
| Cingulum bundle | Cing |
| Stria medullaris thalami | SM |

| Anatomical structure | Abbreviation |
|---|---|
| Medial forebrain bundle, inferior component | MFB (inf.) |
| Ventral tegmental area projection pathway | VTA-PP |
| Optic tract | OT |
| **Internal capsule and commissural pathways** | |
| Anterior limb of the internal capsule | ALIC |
| Posterior limb of the internal capsule | PLIC |
| Retrolenticular part of the internal capsule | RLIC |
| Corpus callosum | CC |
| Forceps major | Fmaj |
| Forceps minor | Fmin |
| Anterior commissure | AC |
| Posterior commissure | PC |
